# Probing Conformational Landscapes of Binding and Allostery in the SARS-CoV-2 Omicron Variant Complexes Using Microsecond Atomistic Simulations and Perturbation-Based Profiling Approaches: Hidden Role of Omicron Mutations as Modulators of Allosteric Signaling and Epistatic Relationships

**DOI:** 10.1101/2023.05.03.539337

**Authors:** Gennady Verkhivker, Mohammed Alshahrani, Grace Gupta, Sian Xiao, Peng Tao

## Abstract

In this study, we systematically examine the conformational dynamics, binding and allosteric communications in the Omicron BA.1, BA.2, BA.3 and BA.4/BA.5 complexes with the ACE2 host receptor using molecular dynamics simulations and perturbation-based network profiling approaches. Microsecond atomistic simulations provided a detailed characterization of the conformational landscapes and revealed the increased thermodynamic stabilization of the BA.2 variant which is contrasted with the BA.4/BA.5 variants inducing a significant mobility of the complexes. Using ensemble-based mutational scanning of binding interactions, we identified binding affinity and structural stability hotspots in the Omicron complexes. Perturbation response scanning and network-based mutational profiling approaches probed the effect of the Omicron variants on allosteric communications. The results of this analysis revealed specific roles of Omicron mutations as “plastic and evolutionary adaptable” modulators of binding and allostery which are coupled to the major regulatory positions through interaction networks. Through perturbation network scanning of allosteric residue potentials in the Omicron variant complexes, which is performed in the background of the original strain, we identified that the key Omicron binding affinity hotspots N501Y and Q498R could mediate allosteric interactions and epistatic couplings. Our results suggested that the synergistic role of these hotspots in controlling stability, binding and allostery can enable for compensatory balance of fitness tradeoffs with conformationally and evolutionary adaptable immune-escape Omicron mutations. Through integrative computational approaches, this study provides a systematic analysis of the effects of Omicron mutations on thermodynamics, binding and allosteric signaling in the complexes with ACE2 receptor. The findings support a mechanism in which Omicron mutations can evolve to balance thermodynamic stability and conformational adaptability in order to ensure proper tradeoff between stability, binding and immune escape.

## Introduction

The SARS-CoV-2 spike glycoprotein is a key molecule involved in the virus’s entry into host cells. It is composed of two subunits, S1 and S2, and is responsible for binding to the host cell surface and for mediating fusion of the viral and cellular membranes. Structural and biochemical studies have revealed the importance of the S glycoprotein in the infection process by demonstrating how the spike protein undergoes stochastic movements between distinct functional forms.^1–9^ The S protein of SARS-CoV-2 has a complex architecture, consisting of an amino (N)-terminal S1 subunit and a structurally rigid carboxyl (C)-terminal S2 subunit. The N-terminal subunit experiences functional motions, with conformational transformations between the closed and open S states. These transitions are enabled by coordinated global movements of the S1 subunit, which is comprised of the N-terminal domain (NTD), the receptor-binding domain (RBD), as well as two structurally conserved subdomains, SD1 and SD2. These components together determine the structural and dynamic response of the S protein to binding partners and the host cell receptor ACE2.^10–15^ The biophysical studies have extensively characterized the thermodynamic and kinetic properties of the SARS-CoV-2 S trimer. In particular, the studies have highlighted the complex interplay between subdomain movements and long-range interactions which couple the S1 and S2 subunits to modulate the RBD equilibrium and population-shifts between the RBD open (up) and closed (down) conformations. This regulation of the RBD exposure and strength of S-ACE2 binding is essential in determining the binding affinity of the S protein to its binding partners.^16–18^ The abundance of cryo-EM and X-ray structures of SARS-CoV-2 S variants of concern (VOCs) in various functional states and complexes with antibodies has allowed for an in-depth understanding of molecular mechanisms and binding epitopes that underlie the binding affinity of the S protein with different classes of antibodies. Structures of the S protein in its various forms have helped to identify conserved and variable regions that are important for antibody binding. The different structures also revealed key residues and structural features that are responsible for the binding of different classes of antibodies to the S protein. This knowledge has been instrumental in the development of effective antibodies and vaccine candidates against SARS-CoV-2.^19–28^ The cryo-EM structures of the S Omicron BA.1 variants in different functional states reveal the intricate balance and tradeoffs of various factors driving binding thermodynamics. It was suggested that Omicron mutations act cooperatively to regulate the open-closed equilibrium, promoting the formation of RBD-up states that induce immune evasion by altering antibody epitopes, while simultaneously maintaining a structurally stable RBD-down state that allows for the occlusion of highly immunogenic sites. This delicate balance is essential for the virus to successfully evade the host immune response.^27^ The Omicron sites N856K, N969K, and T547K have been shown to be important for the stability of the S Omicron BA.1 trimer. Cryo-EM structures have demonstrated that these sites can promote favorable electrostatic interactions and lead to the formation of hydrogen bonds with D658, Q755, and S982 from neighboring subunits, thus increasing the number of inter-protomer contacts. The increased stability of these interactions and the increased number of the inter-protomer contacts which can confer the enhanced stability for the Omicron S-trimer.^28^ The biophysical analysis of protein stability for the Wu-Hu-1, Delta, and Omicron variants was also done by using differential scanning fluorimetry (DSF) assay. The DSF assay measured the inflection temperature, which revealed that the transition to the folding state for the S Omicron BA.1 variant was shifted to lower temperatures compared to the S Wu-Hu-1 and S Delta, suggesting a reduced protein stability of the S Omicron BA.1 variant.^29, 30^ The thermostability of the S-D614G, S-BA.1, and SB-BA.2 protein ectodomains was evaluated in DSF assays and it was found that the BA.1 RBD had reduced stability, while the BA.2 RBD was found to be more stable than BA.1 but less stable than the Wu-Hu-1 protein.^31, 32^ The surprising discovery of structural studies was that the Omicron BA.1 S trimer is likely to adopt the 1RBD up conformation both before and after ACE2 binding. It was found that the S371L, S373P and S375F substitutions generally stabilize the 1RBD-up conformation and prevent exposure of more up-RBDs to ACE2 binding.^33^ Other structural investigations on the Omicron BA.1 variant have also yielded similar findings, suggesting that mutations in the Omicron strain induce enhanced inter-domain and inter-subunit packing that stabilize the open S conformation.^34, 35^ Structural studies provided insight into the evolutionary forces driving the development of the Omicron variant by elucidating the thermodynamic factors that influence the interplay between the ACE2 binding affinity and immune escape.^36–40^ These investigations suggested that mutations increasing ACE2 affinity are balanced with mutations that disfavor binding to ACE2 but facilitate immune escape. The cryo-EM study of the S Omicron BA.1 variant examined binding and antigenic properties by bio-layer interferometry showing that the Omicron variant contains a combination of both types of mutations, with T478K, Q493R, G496S, and Q498R enhancing ACE2 binding while K417N and E484A can decrease the affinity.^41^ This highlights the importance of understanding the thermodynamic factors that drive virus evolution, as these insights can potentially provide new strategies for controlling the virus and preventing its spread. Atomic force microscopy (AFM) studies have consistently demonstrated the importance of mechanical stability in mediating immune evasion, suggesting that a combination of mechanical forces, protein stability, and binding interactions play an important role in controlling the virus’s fitness advantage and immune escape mechanisms.^42–44^

The Omicron BA.2 subvariants of SARS-CoV-2 have been associated with increased transmissibility, severity of disease, and possible vaccine evasion capacity. The binding affinities of Omicron BA.1.1 and BA.2 with ACE2 were found to be stronger than those of BA.3 and BA.1, according to the structures of the RBD-ACE2 complexes for the BA.1.1, BA.2, and BA.3 variants.^45^ The Omicron BA.2 trimer showed higher ACE2 binding affinity compared to both the S Wu-Hu-1 trimer and the S Omicron BA.1 trimer. Specifically, the binding affinity of the Omicron BA.2 trimer was 11 times higher than that of the S Wu-Hu-1 trimer and 2 times higher than that of the S Omicron BA.1 trimer.^46^ The surface plasmon resonance (SPR) results showed that the Omicron BA.4/5 RBD had only a slightly higher binding affinity for ACE2 than the ancestral Wu-Hu-1 strain and the BA.1 variants.^47^ The cryo-EM structures and biochemical analysis of the S trimers for BA.1, BA.2, BA.3, and BA.4/BA.5 subvariants revealed that the BA.2 variant has a higher binding affinity than the other variants, while the BA.4/BA.5 variants have decreased binding affinity.^48^ Structural studies of the Omicron BA.1, BA.2, BA.2.12.1, BA.4, and BA.5 subvariants showed that the BA.2 subvariants had an increased ACE2 binding affinity, a stronger evasion of neutralizing antibody responses compared to the Wu-Hu-1 and Delta strains, and an increased sensitivity towards neutralizing antibodies compared to the other Omicron sublineages.^49^

These findings suggested that the improved ACE2 receptor binding and stronger immune evasion capacity of the Omicron BA.2 subvariants may have contributed to the rapid spread of these variants, as well as their potential to cause more severe disease and possible vaccine evasion.

The structural-functional studies of the Omicron BA.1, BA.2, BA.2.12.1, BA.4, and BA.5 subvariants showed that the BA.2 subvariants had an increased ACE2 binding affinity and a stronger evasion of neutralizing antibody responses compared to the Wu-Hu-1 and Delta strains. This finding confirms that the combined effect of the enhanced ACE2 receptor binding and stronger immune evasion may have contributed to the rapid spread of these Omicron sublineages.^49^ The structure of the BA.4/5 variant of the S protein has been studied in-depth, revealing a number of residue mutations that increase the protein’s affinity for both human ACE2 (hACE2) and mouse ACE2 (mACE2). Residues N501Y, Q493R, G496S, and Q498R increase the number of interactions with mACE2, while the F486V mutation results in the loss of hydrophobic contacts with hACE2 residues F28 and Y83. The net effect of these mutations is that BA.4/5 maintains high affinity interactions with both hACE2 and mACE2, allowing it to balance the need for immune evasion with the need for high-affinity receptor binding.^50^ Another study examined how Omicron BA.4/BA.5 S variants could affect its resistance to neutralizing antibodies, as well as its binding affinity to ACE2, showing that a reversion of R493Q could potentially contribute to attenuating the resistance to neutralizing antibodies, which was rescued by an F486V substitution.^51^ Additionally, an L452R substitution was found to compensate for the decreased ACE2 binding affinity. These results suggest that certain variant substitutions can influence the ability of the S protein to resist neutralizing antibodies and bind to ACE2. Various studies further confirmed that the reversal R493Q in BA.4/BA.5 together with F486V/L452R increased immune evasion capability while retaining binding affinity as compared to the Omicron BA.1 variant.^52–54^ Structural studies of the RBD binding with mouse ACE2 discovered that the Omicron RBD is adapted to mACE2 better than to hACE2 with mutations Q493R, Q498R and Y505H providing stronger interactions to mACE2, while the N501Y mutation is adapted to both hACE2 and mACE2.^55^ These studies suggested that the acquisition of functionally balanced substitutions, where some mutations enhance immune evasion but tend to reduce ACE2 affinity, while others increase ACE2 affinity to compensate for the immune resistant modifications, may be a common strategy of SARS-CoV-2 evolution shared by Omicron subvariants. Overall, the structural-functional studies of the Omicron sublineages suggest that the combined effect of increased ACE2 binding and immune evasion, increased mutation rate, and increased replication rates likely played a role in their rapid spread.

Computer simulations of the SARS-CoV-2 S proteins have allowed for a better understanding of the molecular mechanisms of viral entry and receptor binding.^56–65^ Through simulations, it was revealed that glycosylation plays an important role in modulating the binding of ACE.^56–58^ Molecular simulations also provided insights into the conformational changes of the S protein in the viral membrane, as well as its interactions with other viral and cellular components.^56–60^ Using computational approaches it was suggested that the S protein could function as an allosteric regulatory machinery controlled by stable allosteric hotspots acting as drives drivers and regulators of spike activity.^66–72^ These hotspots can be used to modulate binding functions, allowing the S protein to dynamically respond to other proteins and coordinate the activities of multiple proteins in order to achieve a desired outcome. The all-atom MD simulations of the S Omicron trimer and the Omicron RBD–ACE2 complexes revealed that the Omicron mutations could have enhanced the virus’s infectivity by improving the RBD opening, increasing the binding affinity with ACE2, and optimizing the capacity for antibody escape. This suggests that the virus has evolved to become more effective at infecting human cells by optimizing these three factors.^73^ MD simulations of the Omicron RBD–ACE2 complexes showed that the K417N and G446S mutations reduce the ACE2 binding affinity due to the introduction of polar residues that disrupt the hydrophobic environment that is necessary for the binding of Omicron RBD and ACE2. The Y505H mutation also reduces the binding affinity because the hydrophobic Y505 residue is replaced with a hydrophilic H505 residue, thus further disrupting the hydrophobic environment. On the other hand, the S447N, Q493R, G496S, Q498R, and N501Y mutations were observed to improve the binding affinity with the ACE2 receptor. This is because these mutations introduce polar residues that form hydrogen bonds and hydrophilic interactions with ACE2, thus increasing the binding affinity.^74, 75^

The dynamics-based network analyses showed that the Omicron variant had more functional hotspots and long-range allosteric communications with ACE2 than the Delta variant, suggesting that the Omicron variant may be more effective in mediating binding interactions and allosteric communications. In comparison, the Delta variant appears to have fewer functional hotspots and less long-range allosteric communications with ACE2, suggesting that it may be less effective at mediating such interactions.^76^ The results of these analyses suggest that the Delta and Omicron variants may have different roles in mediating binding and allosteric communications between ACE2 and the S-RBD complexes, and this could lead to different outcomes in terms of viral escape mutants. By combining MD simulations of the Omicron RBD-ACE2 complexes and a systematic mutational scanning of the RBD-ACE2 binding interfaces, it was found that the key Omicron mutational sites R493, R498 and Y501 play a critical role in the binding and activity of the complex.^77^ These sites act as binding energy hotspots, drivers of electrostatic interactions, and mediators of epistatic effects and long-range communications. The results of this study provide valuable insight into the structural and functional properties of the Omicron subvariants of the RBD-ACE2 complex and can be used to inform future drug discovery and design efforts.^77^

In the current study, we employ a combination of atomistic MD simulations and perturbation profiling approaches to systematically examine the conformational dynamics, binding and allosteric communications in the Omicron BA.1, BA.2, BA.3 and BA.4/BA.5 complexes with ACE2. Microsecond atomistic simulations provide a detailed characterization of the conformational landscapes and revealed the increased thermodynamic stabilization of the BA.2 variant which is contrasted with the BA.4/BA.5 variants inducing a significant mobility of the complexes. Using ensemble-based mutational scanning of binding interactions, we characterized binding affinity and structural stability hotspots in the Omicron complexes. Perturbation response scanning and network-based mutational profiling approaches are introduced to probe the effect of the Omicron variants on allosteric communications and characterize roles of Omicron mutations in modulating dynamics, binding and allostery

Through perturbation network scanning of allosteric residue potentials in the Omicron variant complexes, which is performed in the background of the original strain, we show that the key Omicron binding affinity hotspots N501Y and Q498R mediate allosteric interactions and epistatic couplings. Our results suggested that the synergistic role of these hotspots in controlling stability, binding and allostery can enable for compensatory balance of fitness tradeoffs with conformationally and evolutionary adaptable immune-escape Omicron mutations. Through integrative computational approaches, this study provides a systematic analysis and comparison of the effects of Omicron subvariants BA.1, BA.2, BA.3, and BA.4/BA.5 on conformational dynamics, binding and allosteric signaling in the complexes with the ACE2 receptor.

## Materials and Methods

### Structural modeling and refinement

The crystal structures of the BA.2 RBD-hACE2 (pdb id 7WBP), BA.2 RBD-hACE2 (pdb id 7XB0), BA.3 RBD-hACE2 (pdb id 7XB1), and BA.4/BA.5 RBD-hACE2 complexes (pdb id 7XWA) (Figure 1) were obtained from the Protein Data Bank.^78^ During structure preparation stage, protein residues in the crystal structures were inspected for missing residues and protons. Hydrogen atoms and missing residues were initially added and assigned according to the WHATIF program web interface.^79^ The missing loops in the studied cryo-EM structures of the SARS-CoV-2 S protein were reconstructed and optimized using template-based loop prediction approach ArchPRED.^80^ The side chain rotamers were refined and optimized by SCWRL4 tool.^81^ The protein structures were then optimized using atomic-level energy minimization with composite physics and knowledge-based force fields implemented in the 3Drefine method.^82, 83^

**Figure 1.**
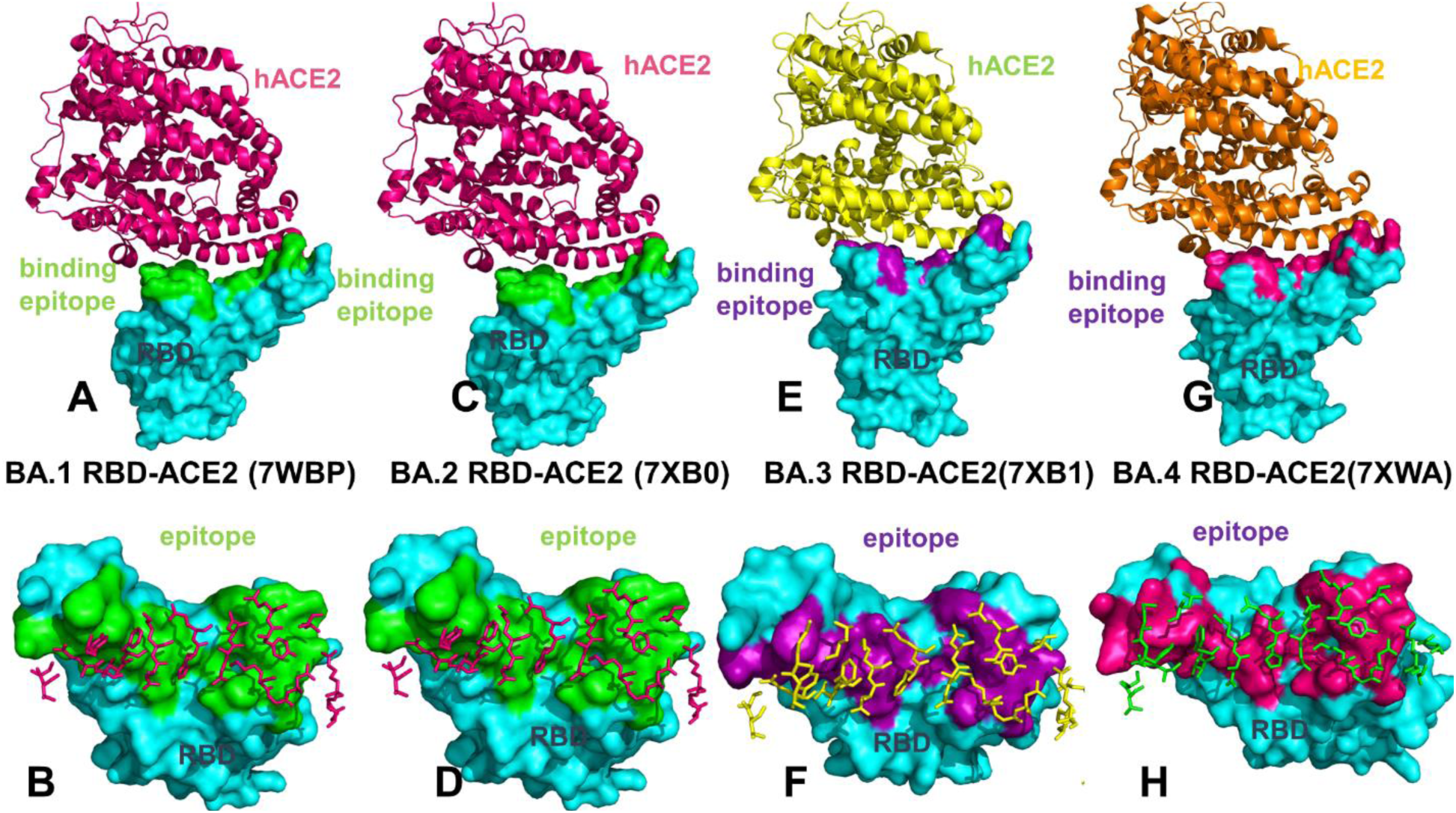
Structural organization and binding epitopes of the SARS-CoV-2-RBD Omicron complexes with human ACE enzyme. (A) The cryo-EM structure of the Omicron RBD BA.1 ACE2 complex (pdb id 7WBP). The RBD is in cyan surface and ACE2 is in pink ribbons. (B) The RBD-BA.1 binding epitope is in green surface. The ACE2 binding residues are shown in pink sticks. (C) The cryo-EM structure of the Omicron RBD BA.2-ACE2 complex (pdb id 7XB0). The RBD is in cyan surface and the ACE2 is in pink ribbons. (D) The RBD-BA.2 binding epitope is shown in green surface. The ACE2 binding residues are shown in pink sticks. (E) The cryo-EM structure of the Omicron RBD BA.3-ACE2 complex (pdb id 7XB1). The RBD is in cyan surface and the ACE2 is in green ribbons. (F) The RBD-BA.2 binding epitope is shown in purple surface and the ACE2 binding residues are shown in green sticks. (G) The cryo-EM structure of the Omicron RBD BA.4/BA.5-ACE2 complex (pdb id 7XWA). The RBD is shown in cyan surface and the ACE2 is in orange ribbons. (H) The RBD BA.4/BA.5 binding epitope is shown in dark-pink surface and the ACE2 binding residues are shown in green sticks.

### All-Atom Molecular Dynamics Simulations

The CHARMM36 force field^84^ with the TIP3P water model^85^ were employed to perform all atom MD simulations for each of the Omicron RBD-hACE2 complexes. The structures of the SARS-CoV-2 S-RBD complexes were prepared in Visual Molecular Dynamics (VMD 1.9.3).^86^ The protonation states of titratable residues were determined under neutral pH. The protein systems were solvated in 130 Å × 85 Å × 75 Å water boxes. In each system, sodium and chloride ions were added to maintain an ionic strength of 0.1 M. After energy minimization, the systems were first heated up from 100 to 300 K with a temperature increment of 20 K per 50 picoseconds (ps). Consequently, the systems were subjected to 1.5 nanoseconds (ns) isothermal−isobaric (NPT) equilibrations at 300 K (equilibrium run), followed by 1 microsecond (µs) canonical (NVT) simulations (production run) at 300 K. Snapshots of the production run were saved every 100 ps. In all simulations, the SHAKE constraint was used to constrain bonds associated with hydrogen atoms in the solvent molecules and the proteins.^87^ The nonbonding interactions within 10 Å were calculated explicitly. The Lennard-Jones interactions were smoothed out to zero at 12 Å. The long-range electrostatic interactions were calculated using the particle mesh Ewald method^88^ with a cut-off of 1.0 nm and a fourth order (cubic) interpolation. The simulations were conducted using OpenMM (version 7.6.0).^89^ For each system, MD simulations were conducted three times in parallel to obtain comprehensive sampling. Each individual simulation of these 5 systems has 10,000 frames. We have 150,000 frames for all simulations in total.

### Mutational Scanning and Sensitivity Analysis

We conducted mutational scanning analysis of the binding epitope residues for the SARS-CoV-2 RBD-ACE2 complexes. Each binding epitope residue was systematically mutated using all substitutions and corresponding protein stability and binding free energy changes were computed. BeAtMuSiC approach^90–92^ was employed that is based on statistical potentials describing the pairwise inter-residue distances, backbone torsion angles and solvent accessibilities, and considers the effect of the mutation on the strength of the interactions at the interface and on the overall stability of the complex. We leveraged rapid calculations based on statistical potentials to compute the ensemble-averaged binding free energy changes using equilibrium samples from simulation trajectories. The binding free energy changes were computed by averaging the results over 1,000 equilibrium samples for each of the studied systems.

### Perturbation Response Scanning

Perturbation Response Scanning (PRS) approach^93–95^ follows the protocol originally proposed by Bahar and colleagues^96, 97^ and was described in detail in our previous studies.^98, 99^ In brief, through monitoring the response to forces on the protein residues, the PRS approach can quantify allosteric couplings and determine the protein response in functional movements. In this approach, it 3N × 3*N* Hessian matrix whose elements represent second derivatives of the potential at the local minimum connect the perturbation forces to the residue displacements. The 3*N*-dimensional vector of node displacements in response to 3*N*-dimensional perturbation force follows Hooke’s law . A perturbation force is applied to one residue at a time, and the response of the protein system is measured by the displacement vector 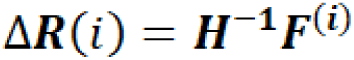 that is then translated into *N*×*N* PRS matrix. The second derivatives matrix is obtained from simulation trajectories for each protein structure, with residues represented by *C*_α_ atoms and the deviation of each residue from an average structure was calculated by 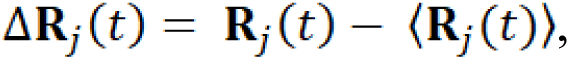, and corresponding covariance matrix C was then calculated by Δ**R**Δ**R***^T^*. We sequentially perturbed each residue in the SARS-CoV-2 spike structures by applying a total of 250 random forces to each residue to mimic a sphere of randomly selected directions. The displacement changes, Δ**R***^i^* is a *3N-*dimensional vector describing the linear response of the protein and deformation of all the residues.

Using the residue displacements upon multiple external force perturbations, we compute the magnitude of the response of residue *k* as 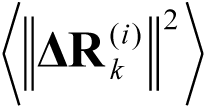 averaged over multiple perturbation forces **F**^(*i*)^, yielding the *ik*^th^ element of the *N*×*N* PRS matrix. The average effect of the perturbed effector site on all other residues is computed by averaging over all sensors (receivers) residues *j* and can be expressed as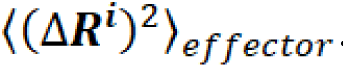. The effector profile determines the global influence of a given residue node on the perturbations in other protein residues and can be used as proxy for detecting allosteric regulatory hotspots in the interaction networks. In turn, the *j* ^th^ column of the PRS matrix describes the sensitivity profile of sensor residue *j* in response to perturbations of all residues and its average is denoted as 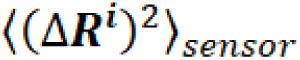. The sensor profile measures the ability of residue *j* to serve as a receiver of dynamic changes in the system.

### Dynamic Network Modeling

A graph-based representation of protein structures^100, 101^ is used to represent residues as network nodes and the inter-residue edges to describe non-covalent residue interactions. The network edges that define residue connectivity are based on non-covalent interactions between residue side-chains. The residue interaction networks were constructed by incorporating the topology-based residue connectivity MD-generated maps of residues cross-correlations^102^ and coevolutionary couplings between residues measured by the mutual information scores.^103^ The edge lengths in the network are obtained using the generalized correlation coefficients associated with the dynamic correlation and mutual information shared by each pair of residues. The length (i.e., weight) of the edge that connects nodes *i* and *j* is defined as the element of a matrix measuring the generalized correlation coefficient as between residue fluctuations in structural and coevolutionary dimensions. Network edges were weighted for residue pairs with in at least one independent simulation. The matrix of communication distances is obtained using generalized correlation between composite variables describing both dynamic positions of residues and coevolutionary mutual information between residues. Residue Interaction Network Generator (RING) program^104, 105^ was employed for generation of the residue interaction networks using the conformational ensemble where edges have an associated weight reflecting the frequency in which the interaction present in the conformational ensemble. The residue interaction network files in xml format were obtained for all structures using RING v3.0 webserver.^106^ Network graph calculations were performed using the python package NetworkX.^107^ Using the constructed protein structure networks, we computed the residue-based short path betweenness parameter. The short path betweenness of residue *i* is defined to be the sum of the fraction of shortest paths between all pairs of residues that pass through residue *i*:

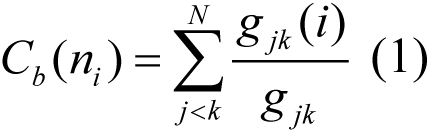

where g*_jk_* denotes the number of shortest geodesics paths connecting *j* and *k*, and *g _jk_* (*i*) is the number of shortest paths between residues *j* and *k* passing through the node *n_i_ .* Residues with high occurrence in the shortest paths connecting all residue pairs have a higher betweenness values. For each node *n*, the betweenness value is normalized by the number of node pairs excluding *n* given as (*N* -1)(*N* -2) / 2, where *N* is the total number of nodes in the connected component that node *n* belongs to. The normalized short path betweenness of residue *i* can be expressed as :

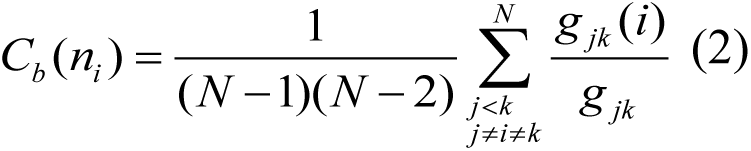

g*_jk_* is the number of shortest paths between residues *j* and k; *g _jk_* (*i*) is the fraction of these shortest paths that pass through residue *i* .

### Network-Based Mutational Profiling of Allosteric Residue Potentials and Epistasis

Through mutation-based perturbations of protein residues we compute dynamic couplings of residues and changes in the short path betweenness centrality (SPC), and the average short path length (ASPL) averaged over all possible modifications in a given position. The change of SPC or ASPL upon mutational changes of each node is reminiscent to the calculation of residue centralities by systematically removing nodes from the network.

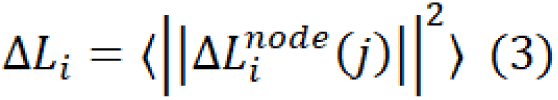

where *i* is a given site, *j* is a mutation and〈⋯〉denotes averaging over mutations. 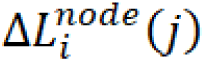 describes the change of SPC or ASPL parameters upon mutation *j* in a residue node Δ*L_i_* is the average change of ASPL triggered by mutational changes in position *i*.

Z-score is then calculated for each node as follows^108, 109^:

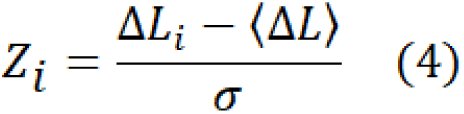

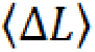 is the change of the SPC or ASPL network parameters under mutational scanning averaged over all protein residues in the S-RBD and σ is the corresponding standard deviation. The ensemble-averaged Z score changes are computed from network analysis of the conformational ensembles using 10,000 snapshots of the simulation trajectory. Through this approach, we evaluate the effect of mutations in the RBD residues on long-range allosteric couplings with the other residues in the RBD-ACE2 complex. We used a measurement based on the Jensen-Shannon divergence (JS) for measuring the similarity between the two distributions of mutation-induced ASPL changes in the Omicron variants relative to the original Wu-Hu-1 strain. Given two distributions, *p* and *q*, both with *g* categories, the Kullback-Leibler (KL) divergence is defined as follows:

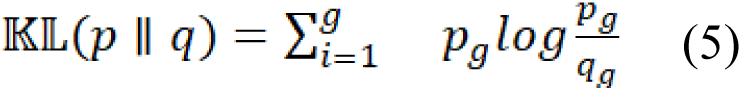

Given two distributions, *p* and *q*, both with *g* categories, the JS divergence is defined as follows:

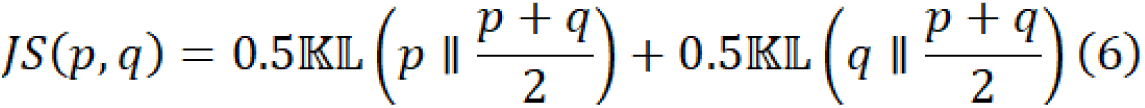

## Results and Discussion

### Microsecond MD Simulations of the Omicron RBD-ACE2 Complexes Reveal Distinct Dynamic Signatures of Stable BA.2 RBD and Highly Mobile BA.4/BA.5 RBD Proteins

Mutations G339D, S373P, S375F, K417N, N440K, S477N, T478K, E484A, Q493R, Q498R, N501Y, and Y505H in BA.2 are shared with the BA.1 variant, but BA.2 additionally carries S371F, T376A, D405N, and R408S mutations (Table 1, Figure 1). G446S mutation is shared with Omicron BA.1, and R493Q reversed mutation is present in BA.4/BA.5 as well as in XBB.1 and XBB.1.5 subvariants. Mutations in F486 are of particular interest as F486V (BA.4/BA.5), F486I, F486S (XBB.1), F486P(XBB.1.5) have been seen in other variants (Table 1) and arguably represent a convergent evolutionary hotspot shared by the recent wave of Omicron subvariants. Structural analysis of the RBD complexes with ACE2 for BA.1 (Figure 1A,B) BA.2 (Figure 1C,D) BA.3 (Figure 1E,F) and BA.4/BA.5 complexes (Figure 1G,H) revealed highly similar RBD conformations, the same binding mode of interactions with the host receptor and virtually identical topography of the binding interface.

**Table 1.** Mutational landscape of the Omicron subvariants in the S-RBD.

| **SARS-CoV-2 Omicron Variant** | **Mutational landscape of the RBD** |
| --- | --- |
| BA.1 | G339D, S371L, S373P, S375F, K417N, N440K, G446S, S477N, T478K, E484A, Q493R, G496S, Q498R, N501Y, Y505H |
| BA.2 | G339D, S371F, S373P, S375F, T376A, D405N, R408S, K417N, N440K, S477N, T478K, E484A, Q493R, Q498R, N501Y, Y505H |
| BA.3 | G339D, S371F, S373P, S375F, D405N, K417N, N440K, G446S, S477N, T478K, E484A, Q493R, Q498R, N501Y, Y505H |
| BA.4/BA.5 | G339D, S371F, S373P, S375F, T376A, D405N, R408S, K417N, N440K, L452R, S477N, T478K, E484A, F486V, R493Q reversal, Q498R, N501Y, Y505H |

**Table 2.** Structures of the Omicron RBD-hACE2 complexes examined in this study.

| **PDB** | **System** | **Per simulation** | **# simulations** |
| --- | --- | --- | --- |
| 7WBP | Omicron RBD BA.1-hACE2 | 1μs | 3 |
| 7XB0 | Omicron RBD BA.2-hACE2 | 1μs | 3 |
| 7XB1 | Omicron RBD BA.3-hACE2 | 1μs | 3 |
| 7XWA | Omicron RBD BA.4/BA.5-hACE2 | 1μs | 3 |

To characterize conformational landscapes and dynamic signatures of the Omicron variants, we conducted several independent microsecond MD simulations of the RBD-ACE2 complexes (Figures 2,3) MD simulations revealed important commonalities and striking differences in the intrinsic conformational dynamics of the RBDs among Omicron variants. Despite structural similarities between the RBD-ACE2 complexes, we found that the Omicron mutations may lead to distinct dynamic profiles in the BA.2 and BA.4/BA.5 RBDs (Figure 2). Moreover, we found Omicron mutations affect conformational dynamics not only through locally induced changes but also induce distinct protein responses over long-range (Figure 2).

**Figure 2.**
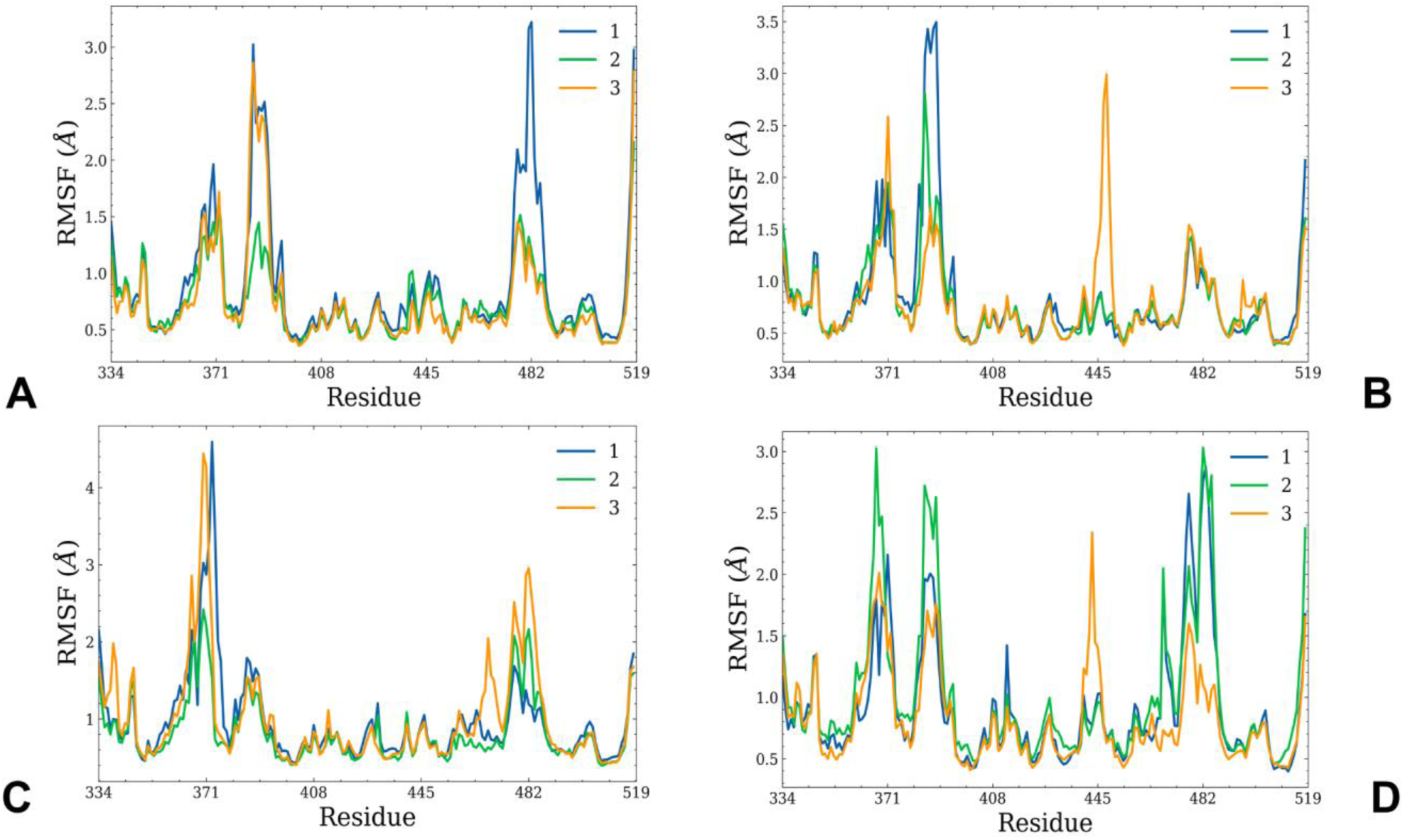
Conformational dynamics profiles obtained from all-atom MD simulations of the Omicron RBD BA.1, BA.2, BA.3 and BA.4/BA.5 complexes with hACE2. The RMSF profiles for the RBD residues obtained from 3 microsecond MD simulations of the Omicron RBD BA.1 hACE2 complex, pdb id 7WBP (A), Omicron RBD BA.2-hACE2 complex, pdb id 7XB0 (B), Omicron RBD BA.3-hACE2 complex, pdb id 7XB1 (C) and Omicron RBD BA.4/BA.5-hACE2 complex, pdb id 7XWA (D).

**Figure 3.**
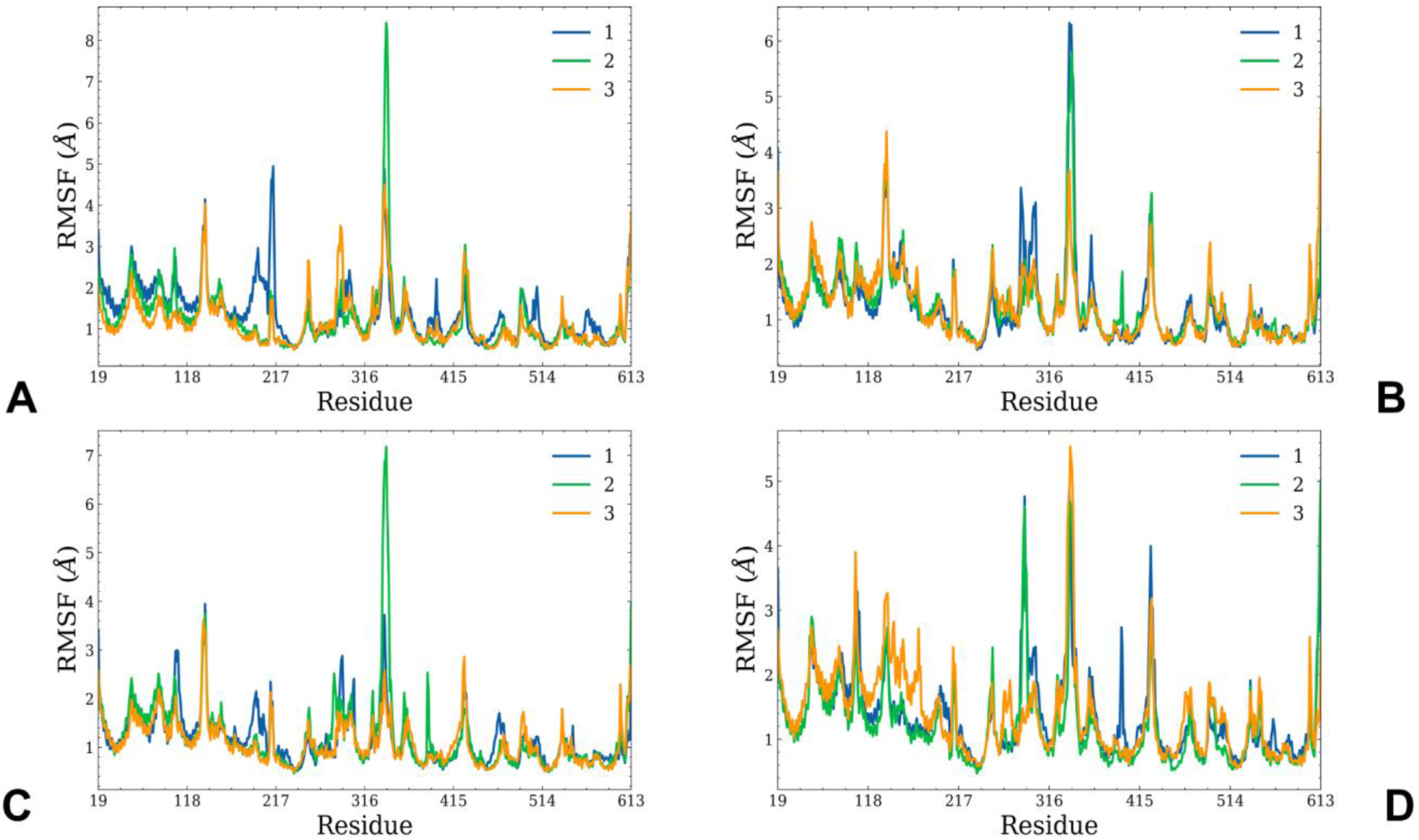
Conformational dynamics profiles of the ACE2 residues obtained from MD simulations of the Omicron RBD BA.1, BA.2, BA.3 and BA.4/BA.5 complexes with hACE2. The RMSF profiles for the ACE2 residues obtained from 3 microsecond MD simulations of the Omicron RBD BA.1-hACE2 complex, pdb id 7WBP (A), Omicron RBD BA.2-hACE2 complex, pdb id 7XB0 (B), Omicron RBD BA.3-hACE2 complex, pdb id 7XB1 (C) and Omicron RBD BA.4/BA.5-hACE2 complex, pdb id 7XWA (D).

The conformational flexibility of the Omicron RBD was analyzed by calculating the root mean square deviations (RMSD) (Supporting Figures S1, S2) and the root mean square fluctuations (RMSF) distribution for the RBD residues (Figure 2A). The RMSD profiles for the RBD residues showed a relatively fast convergence of the MD trajectories, particularly for the BA.2 and BA.3 complexes where all three trajectories converged reaching a steady equilibrium state after about 500 ns (Supporting Figure S1B,C). A somewhat different character of the MD trajectories was seen for BA.1 and BA.4/BA.5 complexes (Supporting Figure S1A,D) reflecting a more dynamic and heterogeneous ensemble of the RBD conformations. The RMSD evolution of the ACE2 residues similarly showed more heterogeneity and greater departures from the crystallographic conformations in the BA.1 and BA.4/BA.5 complexes, while smaller deviations from the experimental structures in more stable BA.2 and BA.3 complexes (Supporting Figure S2). The RMSF profiles showed several local minima regions corresponding to the structured five-stranded antiparallel β-sheet core region that functions as a stable core scaffold. (residues 350-360, 375-380, 394-403) and the interfacial RBD positions involved in the contacts with the hACE2 receptor (residues 440-456 and 490-505 of the binding interface) (Figure 2A). These regions of high structural stability were also seen in our earlier simulation studies of the RBD Wu-Hu-1 and Omicron complexes^76, 77^, further confirming that these segments remain mostly rigid across all examined RBD complexes with hACE2. The dynamics analysis showed that the conformational mobility profiles displayed similar shape reflecting virtually identical structures of the RBD-ACE2 complexes, including the presence of two RMSF peaks corresponding to the flexible peripheral RBD regions (residues 360-373 and residues 380-396). Noticeably, these flexible RBD regions are immediately adjacent to several Omicron mutational sites S371F, S373P, S375F that may partly rigidify the local vicinity of these sites (Figure 2). However, the most striking observation is the differential mobility of the flexible RBM regions (residues 475-490) which ranges from marked stabilization of these residues in the BA.2 RBD (Figure 2B) to the increased mobility in the BA.4/BA.5 variants (Figure 2D).

In all complexes, the conformational dynamics profiles revealed the stability of the hydrophobic core regions including F400, I402, F490, Y453, L455, A475, and Y489 residues (Figure 2). These residues form a network of hydrophobic interactions that play an important role in the recognition and binding of ACE2 receptors. Conformational fluctuations of the RBD core and interfacial RBD residues were significantly restricted in the Omicron complexes.

The trajectories also highlighted the stability of important intermolecular interactions including hydrogen bonds formed by S19 of hACE2 with A475 and N477 of RBD as well as Q24 of hACE2 with N487 of RBD (Supporting Tables S1, S2, Figures 2,3). In addition, Y83 of hACE2 interacts with Y489 and N487 of RBD via hydrogen bonding through π-π stacking interaction with F486 of RBD. Among important stabilizing interactions preserved in simulations are salt bridges between E35 of hACE2 and R493 from RBD in the BA.1, BA.2 and BA.3 complexes. The stability of these interfacial contacts in simulations are consistent with the key role of these contacts in the binding affinity of the Omicron RBD-ACE2 complexes shown experimentally.^19^ The important difference in the dynamic signatures was a considerable stabilization of the RBD in the BA.2 complex (Figure 2B) and BA.3 complex (Figure 2C) as opposed to BA.1 and BA.4/BA.5. In the BA.2 complex, all three MD trajectories displayed a striking reduction in the mobility of the intrinsically flexible RBM loop. Only one of the microsecond trajectories exhibited somewhat larger fluctuations (Figure 2B). MD trajectories of the BA.2 RBD-ACE2 complex showed a considerable degree of convergence in the RMSF values that remained relatively small for the RBD core (RMSF < 1.0 Å) as well as the RBM regions (RMSF < 1.5 Å) including the curtailed fluctuations of the Omicron RBM positions S477N, T478K. The flexibility of the RBM residues was also curtailed in the BA.3 complex in which two trajectories yielded RMSF < 2.5 Å for this region (Figure 2C). At the same time, the flexibility of the RBM region (residues 475-490) increased in BA.1 (Figure 2A) and especially BA.4/BA.5 (Figure 2D). In the BA.4/BA.5 RBD-ACE2 complex, the RBD core residues also showed greater mobility, with some additional local mobility peaks seen for the residues 410-420, 440-450, 470-475. The conformational ensembles revealed that the RBM tip in the BA.2 RBD-ACE2 complex is maintained in a relatively stable fold conformation that can be described as a hook-like folded RBD tip and is similar to the crystallographic conformations. Interestingly, in the BA.1 and especially in a more flexible BA.4/BA.5 RBD, the RBD tip becomes more flexible and often moves away from the “hook” conformation to a more dynamic state in which the RBD tip circulates between a variety of partly disordered conformations. We found that a partly disordered RBM tip can be observed in the ensemble of BA.4/BA.5 RBD ACE2 conformations which is reflected in the appreciably increased RMSFs for this system (Figure 2D). The key Omicron mutational positions Q493R, Q498R, N501Y and Y505H that are involved in the critical ACE2 binding interface region are highly constrained by their strong interactions with the receptor and experience only very minor fluctuations across all the studied variants. Our analysis showed that among the RBD/hACE2 complexes, the BA.2 RBD/hACE2 binding interface has the largest number of highly stable intermolecular contacts and hydrogen bonds (Tables S1,S2). The salt bridges, hydrogen bonds and hydrophobic interactions were very stable in the RBD BA.2 complex and displayed the higher occupancy of the favorable contacts as compared to other Omicron RBD-hACE2 complexes (Table S2).

The RMSF analysis of the ACE2 residues showed similar profiles across all the examined variants (Figure 3). The highly stable ACE2 residues corresponding to the rigid core and the binding interface positions centered around K353 and H34 (Figure 3). The key ACE2 binding motifs correspond to an alpha-helix (residues 24-31) and a beta-sheet (residue 350-356) that display rather moderate RMSF values in all complexes. The important polar /charged residue interactions at the interface are formed with ACE2 residues D30, K31, H34 and E35 that all display very small thermal fluctuations. Other ACE2 residues Q24, M82, Y83, D38, Y41, N330, K353 that anchor different parts of the with the RBD-ACE2 binding interface remained stable in all MD trajectories (Figure 3). Notably, the conformational dynamics profiles revealed minor signs of the increased ACE2 heterogeneity in the BA.4/BA.5 complex (Figure 3D), likely reflecting the reciprocal mobility of the RBD in this complex.

To further examine the character of dynamic couplings and quantify correlations between motions of the RBD regions we performed the dynamic cross correlation (DCC) residue analysis and reported the DCC maps for the Omicron RBD-ACE2 complexes (Figure 4).

**Figure 4.**
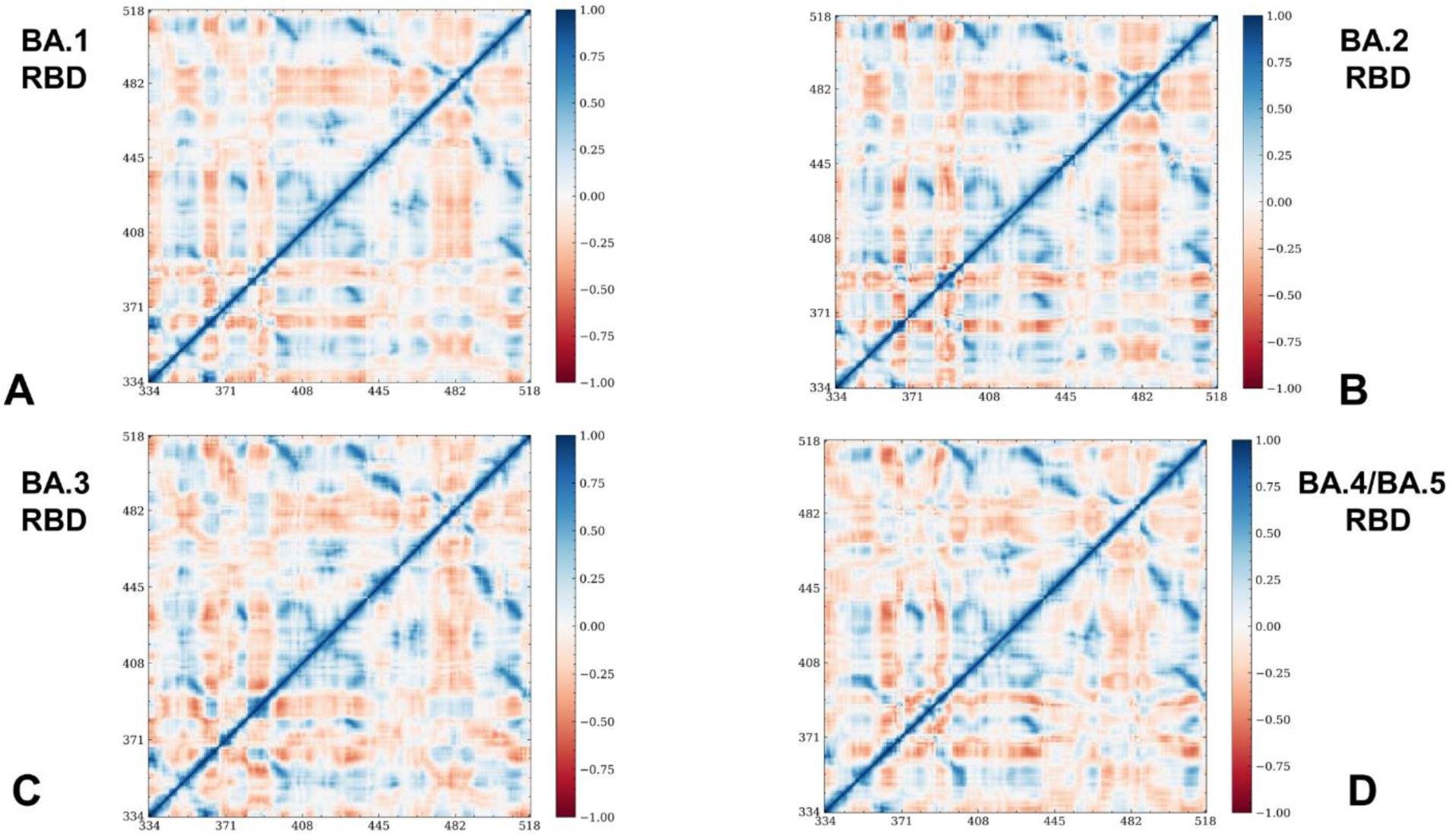
The DCC for the RBD residues in the Omicron RBD BA.1-hACE2 complex, pdb id 7WBP (A), Omicron RBD BA.2-hACE2 complex, pdb id 7XB0 (B), Omicron RBD BA.3 hACE2 complex, pdb id 7XB1 (C) and Omicron RBD BA.4/BA.5-hACE2 complex, pdb id 7XWA (D).

The DCC maps demonstrated subtle but important differences in the dynamic couplings, particularly stronger positive dynamic correlations in the RBD core regions (residues 333-445) for BA.2 (Figure 4B) and BA.3 (Figure 4C). Interestingly, we noticed the presence of negative cross-correlation between motions of the RBM tip (residues 475-485) and other RBD regions (residues 400-470, 490-520) (Figure 4B). This reflects a more stable RBD in the BA.2 complex yielding strong dynamic couplings between the RBD core and various parts of the binding interface. A weaker but similar pattern of the inter-correlated motions was also seen in the BA.1 and BA.3 complexes (Figure 4A,C). Consistent with the conformational dynamics analysis, we found that the correlated motions become appreciably weaker in the BA.4/BA.5 RBD-ACE2 complex (Figure 4D), reflecting the elevated level of mobility in the intrinsically flexible RBM region and the increased thermal fluctuations in the RBD core.

To summarize, by performing multiple microsecond MD simulations of the Omicron RBD ACE2 complexes, we found evidence of distinct dynamic patterns in the structurally similar complex conformations. Consistent with the experimental data, our results showed that BA.2 mutations may induce the increased stabilization of the RBD in the complex with ACE2 which may be directly linked with the higher binding affinity than the other variants, including BA.4/BA.5 variants while a considerably greater flexibility may be the important dynamic attribute of the BA.4/BA.5 variants that have the decreased binding affinity.^48^ Furthermore, the recent pioneering experiments demonstrated that BA.4/BA.5 variants escaped both the vaccine-induced and BA.1 infection-induced antibodies more than BA.1 and BA.2 sub lineages.^47^ It is likely that the experimentally observed distinct antigenic profiles for BA.1, BA.2 and BA.5 Omicron sub-variants^110^ may be linked to their distinct dynamic signatures where the increased immune escape of BA.4/BA.5 variant could be enabled by the elevated mobility and conformational plasticity of the BA.4/BA.5 RBD protein.

### Mutational Sensitivity Analysis Identifies Structural Stability and Binding Affinity Hotspots in the Omicron RBD-ACE2 Complexes

Using the conformational ensembles of the Omicron RBD variant complexes we performed a systematic mutational scanning of the interfacial RBD residues. In silico mutational scanning was done using BeAtMuSiC approach.^90–92^ This approach utilizes statistical potentials and allows for robust predictions of the mutation-induced changes of the binding interactions and the stability of the complex. We enhanced BeAtMuSiC approach by invoking ensemble-based computations of the binding free energy changes caused by mutations. In our adaptation of this method, the binding free energy ΔΔG changes were evaluated by averaging the results of computations over 1,000 samples from MD simulation trajectories. To provide a systematic comparison, we constructed mutational heatmaps for the RBD interface residues in each of the studied Omicron RBD-hACE2 complexes (Figures 5,6).

**Figure 5.**
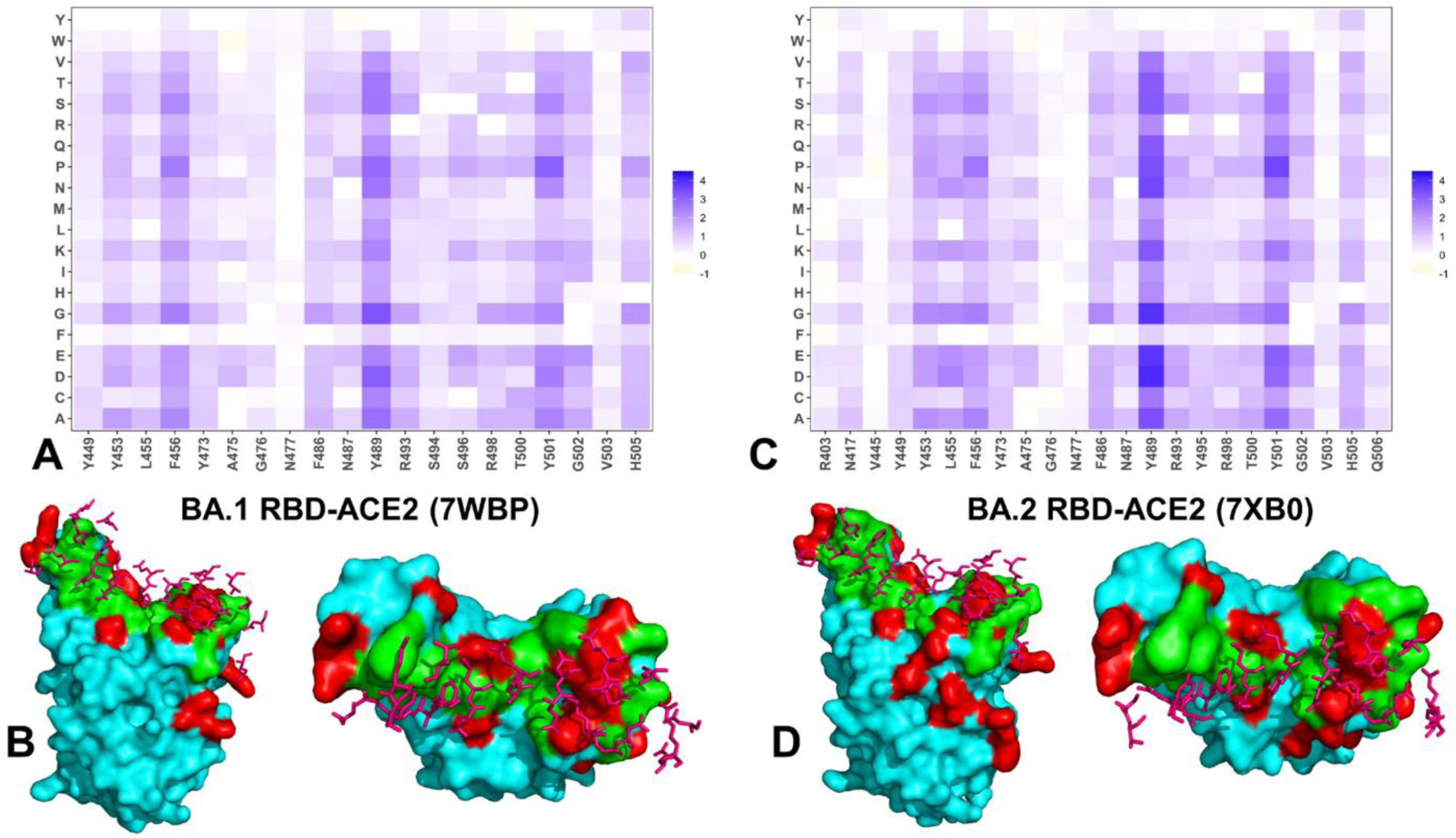
Ensemble-based dynamic mutational profiling of the RBD intermolecular interfaces in the Omicron RBD-hACE2 complexes. The mutational scanning heatmaps are shown for the interfacial RBD residues in the Omicron BA.1 RBD-hACE2 (A) and Omicron BA.2 RBD hACE2 complexes (C). Structural mapping of the RBD binding epitopes of the Omicron BA.1 hACE2 complex (B) and BA.2 RBD-hACE2 (D).The RBD binding epitope is shown in green colored surface. The ACE2 binding residues are in pink sticks. The Omicron RBD BA.1 and BA.2 mutational sites are shown in red surface. The heatmaps show the estimated binding free energy changes for 20 single mutations of the interfacial positions. The standard errors of the mean for binding free energy changes were based on MD trajectories and selected samples (a total of 1,000 samples) are within ∼ 0.07-0.15 kcal/mol using averages from MD trajectories.

**Figure 6.**
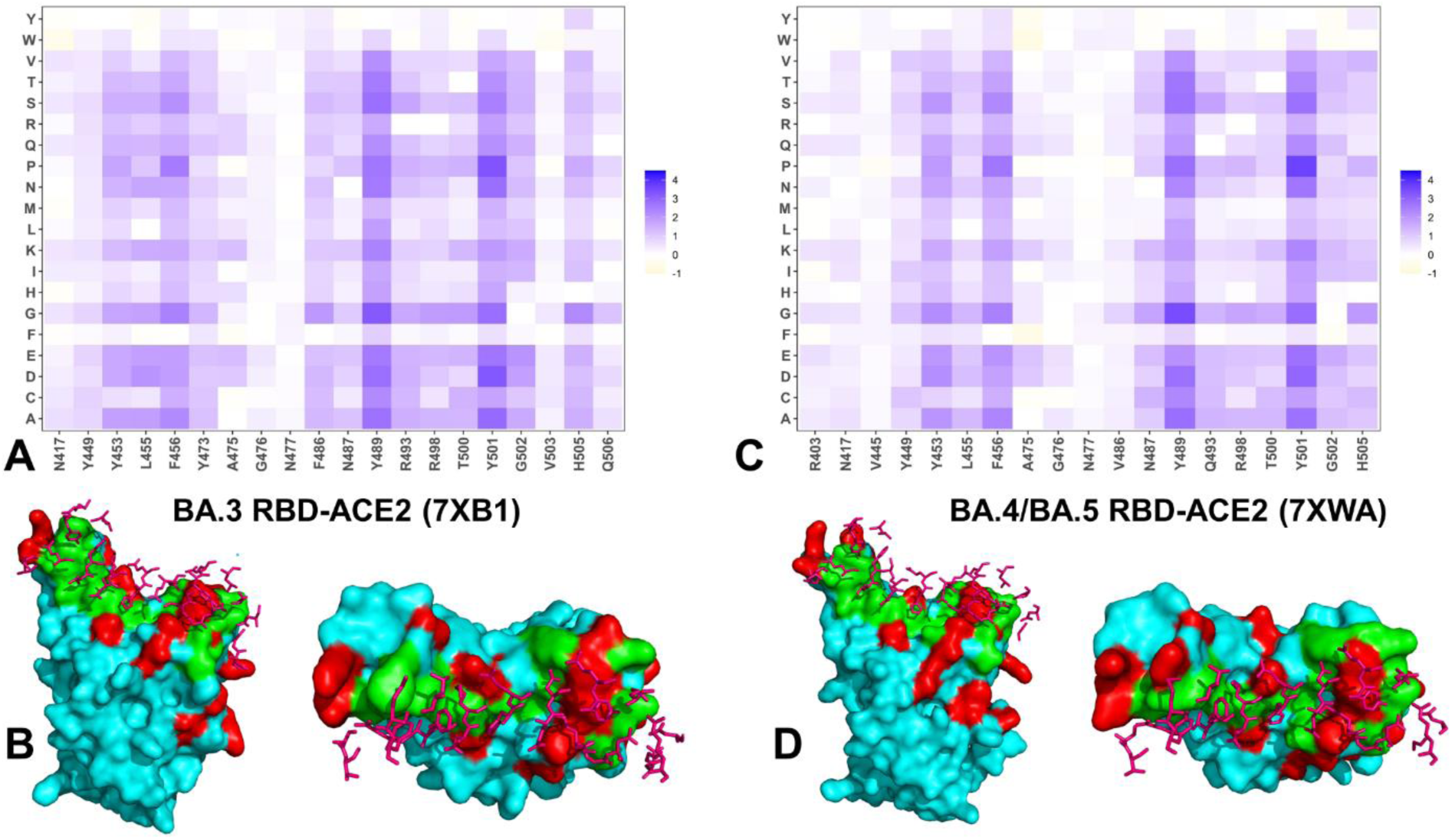
Ensemble-based dynamic mutational profiling of the RBD intermolecular interfaces in the Omicron RBD-hACE2 complexes. The mutational scanning heatmaps are shown for the interfacial RBD residues in the Omicron BA.3 RBD-hACE2 (A) and Omicron BA.4/BA.5 RBD-hACE2 complexes (C). Structural mapping of the RBD binding epitopes of the Omicron BA.3-hACE2 complex (B) and BA.4/BA.5 RBD-hACE2 (D).The RBD binding epitope is shown in green-colored surface. The ACE2 binding residues are in pink sticks. The Omicron RBD BA.3 and BA.4/BA.5 mutational sites are shown in red surface. The heatmaps show the estimated binding free energy changes for 20 single mutations of the interfacial positions. The standard errors of the mean for binding free energy changes were based on MD trajectories and selected samples (a total of 1,000 samples) are within ∼ 0.12-0.18 kcal/mol using averages from MD trajectories.

Consistent with deep mutagenesis experiments^111, 112^ the binding energy hotspots correspond to hydrophobic residues F456, F486, Y489 and Y501 that play a decisive role in binding for Omicron BA.1 and BA.2 complexes (Figure 5). Mutational heatmaps clearly showed that all substitutions in these key interfacial positions can incur a consistent and considerable loss in the stability and binding affinity with ACE2. Mutational scanning of Y453 and L455 also showed significant destabilization changes, particularly for the BA.2 variant (Figure 5B) where both positions belong to the group of major binding hotspot.

Importantly, Y501 and R498 mutational sites are highly favorable for binding, and mutations in these positions induce appreciable and consistent destabilization (Figure 5). At the same time, mutational scanning of R493 generally results in moderate destabilization changes and is less sensitive to perturbations than R498 and Y501 positions. It should be noticed that the Omicron RBD BA.2 complex featured a larger binding interface, in which the mutation-induced destabilization is distributed over more residues (Figure 5B). In addition to the greater RBD stability the stronger binding affinity of the BA.2 complex may also be associated with the larger binding interface and that allows for the greater cumulative contribution of the RBD binding residues. The difference in the RBD-ACE2 interface for the BA.1 and BA.2 variants are the two residues located at residue positions 446 and 496, which are S446 and S496 for BA.1 and G446 and G496 for BA.2. We observed that mutations of S496 is more detrimental for binding. Despite a larger binding interface in BA.2 which featured in addition R403, V445 and G446 residues, these positions are relatively tolerant to perturbations and cause only modest destabilization.

While the binding energy hotspots Y453, L455, F456, Y489, R493, R498, Y501 are shared between BA.2 and BA.3 complexes, the interfacial positions A475, G476 and N477 are more tolerant in the BA.3 (Figure 6A). In addition, positions 502-506 in the BA.3 variant (Figure 6A) are more tolerant to mutations than the respective sites in the BA.2 (Figure 5B). While mutational maps for BA.2 and BA.3 variants are still quite similar, which is consistent with the structural studies and binding affinity measurements, more radical differences were observed in the mutational scanning of BA.4/BA.5 RBD (Figure 6C). Notably, the R403, N417, V445, A475, G476, N477, V486 and N487 sites become much “softer” and more tolerant to substitutions as mutations in these positions induce very minor changes and could be moderately stabilizing. In agreement with deep mutation scans our analysis predicted that the F486V mutation would reduce the affinity of the RBD for hACE2.^111, 112^ The BA.4/5 RBD complex is characterized by loss of hydrophobic contacts for F486V mutation with ACE2 residues F28 and Y83. These changes transpire in weaker binding interactions of F486V position with ACE2 residues leading to more tolerant to substitutions binding free energy changes (Figure 6C,D). At the same time, Q493 can reestablish a hydrogen bond interaction with ACE2 K31 residue, that was lost due to the charge repulsion between R498 and K31. As a result, mutations in Q493 are moderately destabilizing (Figure 6C).

Overall, the observed differences in the mutational scanning map of the BA.4/BA.5 RBD complex showed a broadly distributed weakening of binding interactions across various segment of the binding interface, including strategically important positions R403, N417, V445, A475, G476, N477, V486 and N487 (Figure 6C). At the same time, highly destabilizing mutation induced changes were observed in positions Y453, L455, F456, Y489, Q493, T500, Y501 and H505 (Figure 6C).

Interestingly, our analysis suggested that the L452R mutation may not have a direct impact on the binding interface contacts with ACE2 in the BA.4/BA.5 complex, and the functional role of this Omicron mutation may be manifested through allosteric effects and modulation of antibody escape. These observations are also in line with the body of evidence suggesting that the reversal R49Q combined with F486V/L452R changes in BA.4/BA.5 may enhance the immune escape while maintaining the binding affinity comparable to the Omicron BA.1 variant.^52–54^

### Perturbation Response Scanning Reveals Variant-Specific Modulation of Allosteric Effector and Sensor Centers : Activation of Distinct Allosteric Communication Routes in BA.2 and BA.4/BA.5 Complexes

Using the PRS method, we probed the allosteric effector and sensor potential of the RBD residues in RBD-ACE2 complexes. In this model, the effector profiles evaluate allosteric propensity of protein residues to efficiently propagate signals over long-range in response to systematically applied external perturbations. Accordingly, the local maxima along the effector profile may serve as an indicator of allosteric hotspots that can influence dynamic changes in other residues and may control signal transmission in the system. The effector peaks corresponding to the RBD sites with a high allosteric potential (residues 338-340, 348-353, 400-406, 420-422, 432-436, 450-456, 505-512) are conserved across all RBD-ACE2 complexes (Figure 7A,B) The highest peaks are aligned with the hydrophobic RBD core residues 399-402 and a β7 core RBD segment (residues 506-512) that connects N501Y and Y505H interfacial positions with the central RBD core. The major allosteric effector clusters were also observed in the RBD core (residue 348-353) and functionally important segment 450-456 connecting the central core with the binding interface (Figure 7A,B). Interestingly, the effector cluster peaks were more pronounced for BA.2 and BA.3 RBDs (Figure 7A,B), indicating that the allosteric potential of these RBD residues is enhanced in these variants. Together, these observations suggest that BA.2 and BA.3 RBD may feature a robust network of stable allosteric centers that could mediate an ensemble of well-defined signaling paths from the RBD core to the interface regions connecting to the hotspots near R498/Y501 mutational sites. By mapping positions of the BA.2 mutational sites on the PRS profiles, we noticed that the effector centers are typically not targeted by Omicron mutations, as these hotspots usually correspond to structural stable positions essential for allosteric communication and signal transmission between the RBD and ACE2 proteins. However, stronger allosteric potential was seen for Q493R, Q498R, and especially N501Y and Y505H residues in BA.2 and BA.3 indicating that these positions play a key role in transmitting communication signal between the RBD and ACE2. Our findings suggest that BA.2 mutations may not only induce the increased stabilization of the RBD and enhanced binding interface, but also amplify the allosteric potential of the key binding hotspots, thereby increasing the efficiency of preferential routes for long-range communications with ACE2.

**Figure 7.**
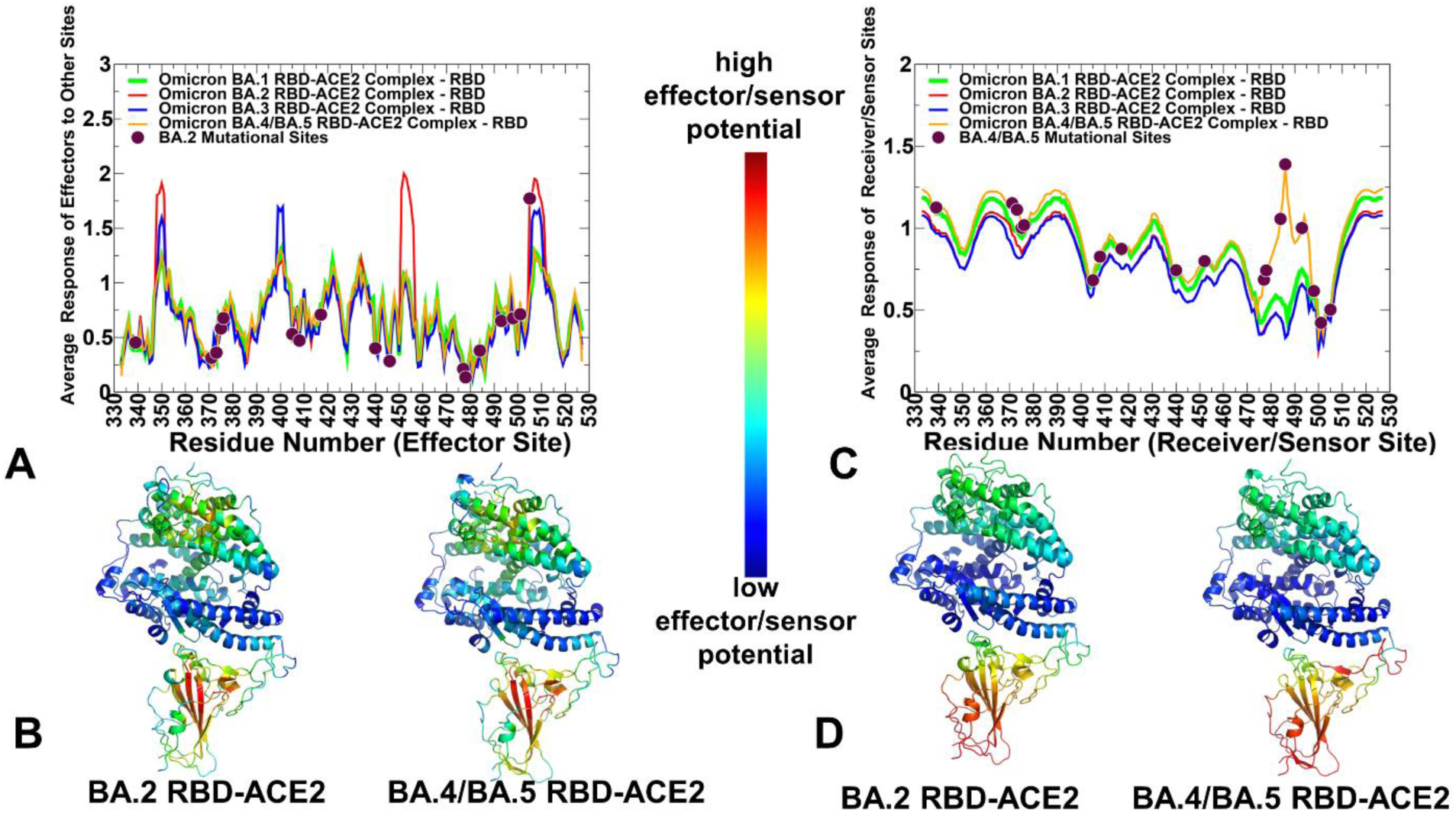
The PRS analysis of the SARS-CoV-2 Omicron RBD-ACE2 (A) The PRS effector profiles for the BA.1 RBD (in green lines), BA.2 RBD (in red lines), BA.3 RBD (in blue lines) and BA.4/BA.5 RBD (in orange lines). The positions of BA.2 RBD mutational sites (G339D, S371F, S373P, S375F, T376A, D405N, R408S, K417N, N440K, S477N, T478K, E484A, Q493R, Q498R, N501Y, Y505H) are indicated by maroon-colored filled circles. (B) Structural maps of the PRS effector profiles are shown for the BA.2 RBD and BA.4/BA.5 RBD. The color gradient from blue to red indicates the increasing effector propensities. (C) The PRS sensor/receiver profiles for the BA.1 RBD (in green lines), BA.2 RBD (in red lines), BA.3 RBD (in blue lines) and BA.4/BA.5 RBD (in orange lines). The positions of BA.4/BA.5 RBD mutational sites (G339D, S371F, S373P, S375F, T376A, D405N, R408S,K417N, N440K, L452R, S477N, T478K, E484A, F486V, Q493, Q498R, N501Y, Y505H) are indicated by maroon-colored filled circles. (D) Structural maps of the PRS sensor profiles are shown for the BA.2 RBD and BA.4/BA.5 RBD.

A more dynamic nature of the BA.1 and especially BA.4/BA.5 complexes yielded the effector profile with a more “diffuse” distribution of relatively weak allosteric mediators (Figure 7C,D). This would imply a less rigid allosteric interaction network in which multiple communication pathways could be activated leading to alternative transmission routes across the binding interface. The sensor profile and respective distribution peaks highlight residues that have a strong propensity to sense signals and produce allosteric response through altered dynamics. The analysis revealed variant-specific modulation of the PRS sensor profiles, in which BA.2 and BA.3 RBDs have similar distributions and small peaks associated with the flexible RBD regions that may transmit the allosteric signals from the regulatory sites (Figure 7C,D).

The analysis suggested that for BA.2 and BA.3 complexes major allosteric communication routes would likely proceed through the central binding hub formed by R498, Y501 and H505 residues. In addition, minor communication routes for these complexes may be offered by the flexible transmission regions (residues 446-460). A different scenario may occur for the BA.4/BA.5 complex in which a strong peak of the PRS sensor profile was associated with the flexible RBM region harboring Omicron mutational sites S477N, T478K, E484A, F486V, and Q493 (Figure 7C,D). As a result, allosteric signal transmission in the BA.4/BA.5 RBD complex may preferentially proceed through the highly flexible and adaptive RBM tip. According to the dynamics analysis, F486V in BA.4/BA.5 could markedly increase the mobility of the RBM flexible loop and sample multiple “disordered” tip conformations as opposed to BA.2 and BA.3 RBDs that preferentially sample the crystallographic state.

These findings suggest that allosteric communications between the RBD and ACE2 in BA.4/BA.5 variant may form a broad ensemble passing through flexible Omicron sites E484A and F486V. Given that F486V can moderately reduce the binding affinity of BA.4/BA.5 RBD with ACE2, this thermodynamic effect may be counterbalanced by the kinetic preferences, providing a possible explanation to the increased transmission of BA.4/BA.5 variants. It is tempting to suggest that the increased mobility and the activation of the transmission potential for the RBM loop residues mediated by E484A/F486V mutations may help to enhance the immune escape fitness. We argue that Omicron mutational sites can be dynamically coupled through short and long-range interactions forming an adaptive allosteric network that controls balance between conformational plasticity, protein stability, and functional adaptability.

### Mutational Profiling of Allosteric Communications in the Omicron RBD-ACE2 Complexes Reveals Hidden Role of Omicron Mutations As Mediators of Allosteric Signaling and Epistatic Couplings

We complemented the PRS results with the network-based mutational profiling of allosteric residue propensities that are computed using topological network parameters SPC and ASPL (see Materials and Methods) that characterize global network of allosteric communications. Through ensemble-based averaging over mutation-induced changes in these network metrics, the proposed model can identify positions in which mutations on average cause network changes. Allosteric hotspots are identified as residues in which mutations incur significant edgetic perturbations of the global residue interaction network that disrupt the network connectivity and cause a significant impairment of global network communications and compromise signaling.

By performing in silico version of “deep” mutational scans to measure the allosteric effects in the RBD-ACE2 complexes, we examine the variant-induced network changes in the background of the original Wu-Hu-1 strain. Using the network-based mutational profiling approach, we can characterize whether Omicron mutations cause synergistic changes in allosteric communications that may emulate potential epistatic couplings in the effects of mutations at other sites.

Using a graph-based network model of residue interactions we computed the ensemble averaged distributions of the residue-based SPC metric (Figure 8). This centrality metric is based on computing of the short path betweenness as outlined in detail in the Methods section. By systematically introducing mutational changes in the RBD, we computed the ensemble averaged mutation-induced changes in the SPC parameter.

**Figure 8.**
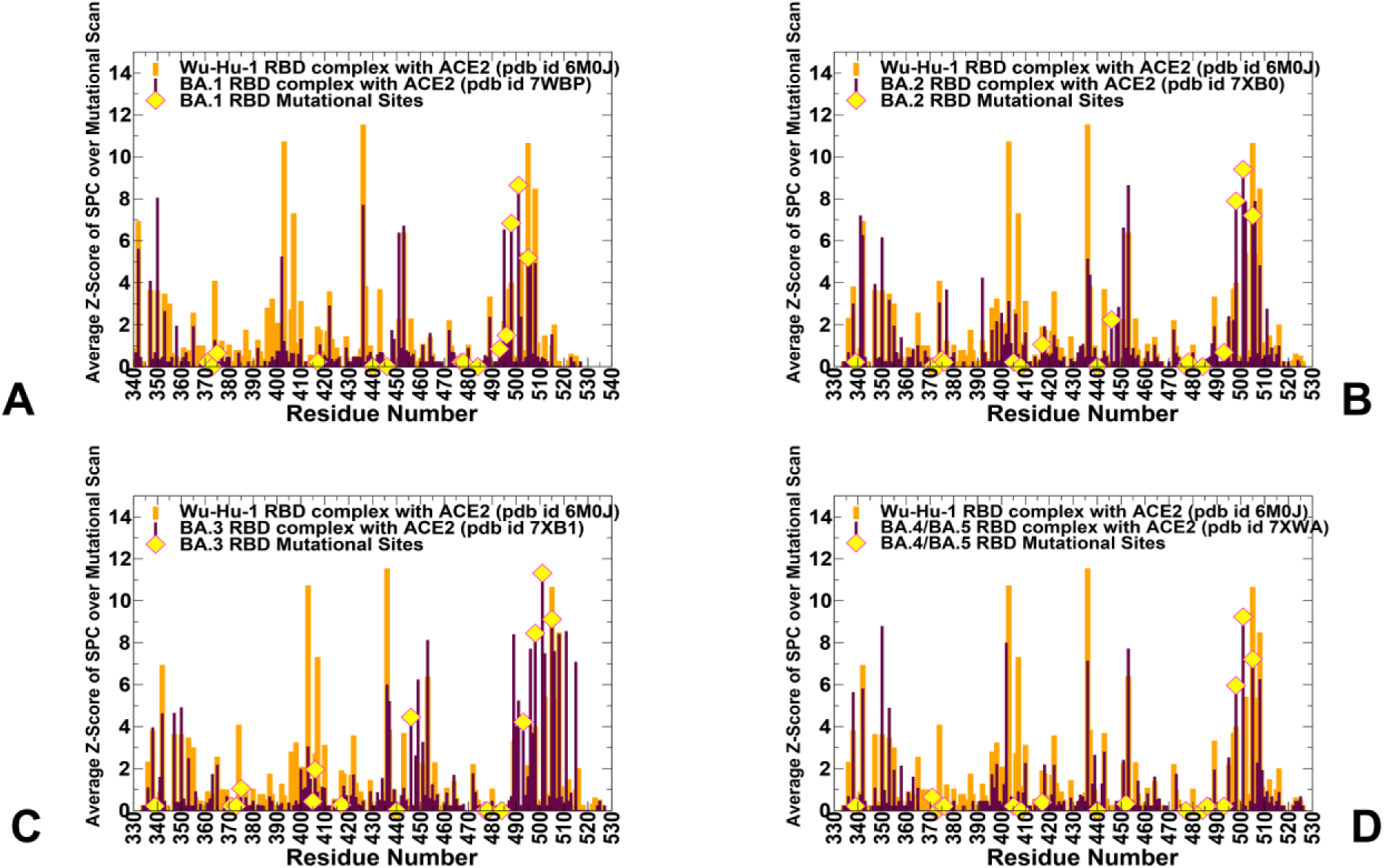
The dynamic network-based analysis of the Omicron RBD subvariant complexes with ACE2 in the background of the original Wu-Hu-1 strain. The Z-score of SPC centrality for RBD residues averaged over mutational scan for the BA.1 RBD-ACE2 complex (A), BA.2 RBD-ACE2 complex (B), BA.3 RBD-ACE2 complex (C) and BA.4/BA.5 RBD-ACE2 complex (D). The distributions for Omicron RBDs are shown in maroon bars and the distribution for the Wu-Hu-1 RBD is shown in orange bars. The positions of the Omicron mutations are highlighted on the distribution profiles in yellow-colored filled circles.

The distributions revealed several critical clusters of residues that are important for mediating allosteric communications, including the RBD core segments (residues 340-355, 400-410, 430-450) and binding interface residues 495-505 (Figure 8). While in the background Wu-Hu-1 variant the distribution is dominated by the RBD core clusters (residues 400-410 and 430-450),

Omicron BA.2 and BA.3 variants can induce changes in the relative contribution of allosteric centers towards the binding interface region anchored by R498, Y501 and H505 positions (Figure 8B,C). Moreover, the profiles for BA.2 and BA.3 variants displayed an extended and dense cluster of allosteric centers (residues 495-510) that links the RBD core residues with the binding interface hotspots R498, Y501 and H505. In both BA.2 and BA.3 profiles, we observed a synergistic increase in allosteric propensities of Omicron mutations R498, Y501 and H505 as compared to the background distribution. This may exemplify the enhanced density of allosteric communication routes in BA.2 passing through the binding hotspots that may act cooperatively and exert potential epistatic effects. In some contrast, the distribution for the BA.4/BA.5 complex was more similar to the Wu-Hu-1 profile (Figure 8D), and yet the peaks associated with R498 and Y501 residues also emerged synchronously and were appreciably larger than the Z-scores for Q498 and N501 in the original strain. In this context, the mutation induced changes in the network distributions for the Omicron variants are similar to the experimentally determined profile of the epistatic shifts dominated by N501Y and Q498R and less significant shifts experienced by residues 446-449 and residues 505, 506. ^113, 114^

Perturbation-based profiling of allosteric residue propensities using mutational scanning of the ASPL changes provided more information about potential allosteric hotspots by mapping a space of network-altering allosteric ‘edgetic’ variant sites (Figures 9,10). In this model, we characterize residues where mutations on average induce a significant increase in the ASPL metric and therefore have a dramatic effect on the efficiency of long-range communications in the allosteric interaction network. This analysis enables identification of allosteric control points that could determine the efficient and robust long-range communications in the complexes. The distributions of the average Z-score of ASPL over mutations are characterized by a group of conserved peaks that are shared across all complexes. The commonly shared allosterically important positions include F338, V341, F342, F347, V350, F377, F392, 400-403, W436, Y451, L452, Y495, Y505H, and Y508 (Figures 9,10). Not surprisingly, a significant fraction of these allosteric centers are stable hydrophobic sites that are strategically located in the RBD core and mediate network of communications between the RBD residues. Another important revelation of this analysis was an appreciable correspondence between the PRS effector centers (residues 348-353, 400-406, 420-422, 432-436, 505-512) and the predicted allosteric centers mediating efficient communications in the complexes.

**Figure 9.**
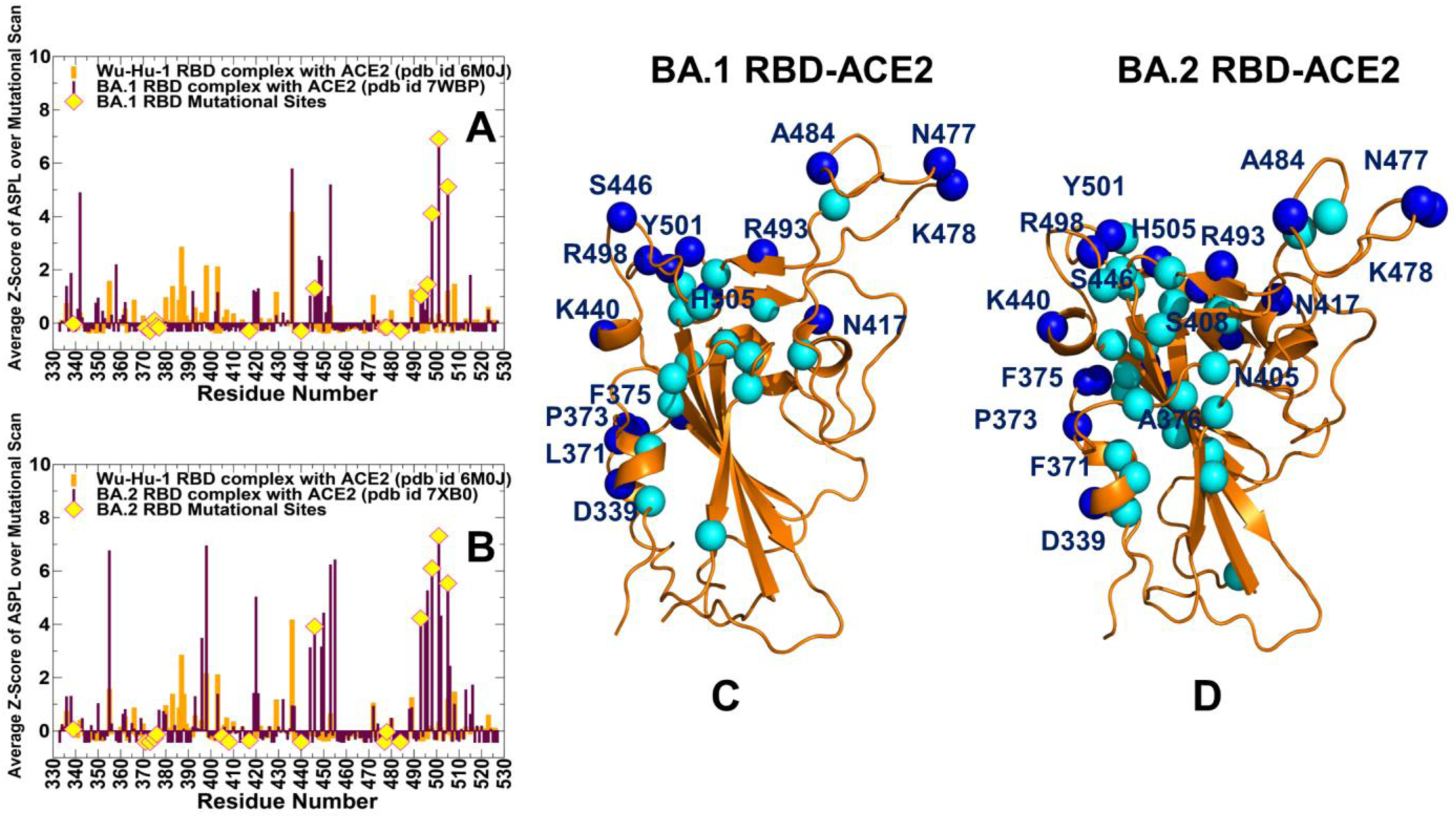
The dynamic network-based analysis of the Omicron RBD subvariant complexes with ACE2 in the background of the original Wu-Hu-1 strain. The Z-score of ASPL for RBD residues averaged over mutational scan for the BA.1 RBD-ACE2 complex (A) and BA.2 RBD-ACE2 complex (B). The distributions for Omicron RBDs are shown in maroon bars and the distribution for the Wu-Hu-1 RBD is shown in orange bars. The positions of the Omicron mutations are highlighted on the distribution profiles in yellow-colored filled circles. Structural mapping of the RBD residues with high allosteric potential (in cyan spheres) in the BA1 RBD (C) and BA.2 RBD (D). The Omicron mutational sites are in blue spheres and annotated.

**Figure 10.**
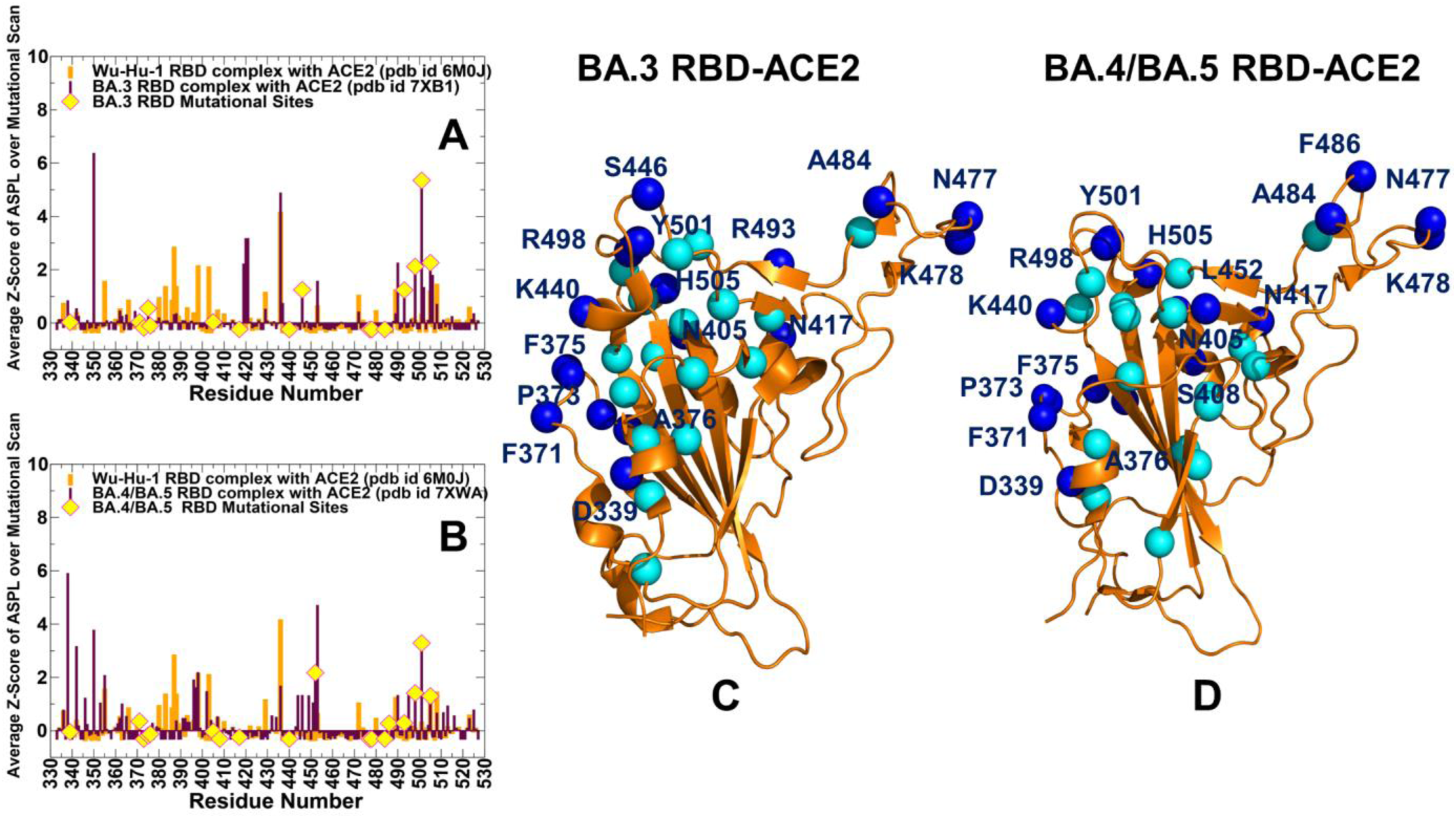
The dynamic network-based analysis of the Omicron RBD subvariant complexes with ACE2 in the background of the original Wu-Hu-1 strain. The Z-score of ASPL for RBD residues averaged over mutational scan for the BA.3 RBD-ACE2 complex (A) and BA.4/BA.5 RBD-ACE2 complex (B). The distributions for Omicron RBDs are shown in maroon bars and the distribution for the Wu-Hu-1 RBD is shown in orange bars. The positions of the Omicron mutations are highlighted on the distribution profiles in yellow-colored filled circles. Structural mapping of the RBD residues with high allosteric potential (in cyan spheres) in the BA.3 RBD (C) and BA.4/BA.5 RBD (D). The Omicron mutational sites are in blue spheres and annotated.

The distributions revealed important differences that can be exemplified by comparison of BA.1 (Figure 9A) and BA.2 variants (Figure 9B) in the background of the original strain. Strikingly, we found that BA.2 variant mutations induced significant redistribution of the Wu-Hu-1 profile, resulting in the emergence of two major clusters of allosteric sites. The major allosteric cluster is fairly broad (residues 495-510) and is anchored by R498, Y501 and H505 BA.2 sites that dominate the distribution (Figure 9B). Notably, Y501 is aligned with the largest peak in this cluster. These observations highlighted the collective emergence of these sites as major allosteric mediating centers in the background of the original strain. In network terms, this implies strong synergistic couplings between R498, Y501 and H505 centers to preferentially direct allosteric communications between the RBD and ACE2 molecules. As our analysis is based on systematic mutational scanning of allosteric residue potentials measuring changes in the average short paths, the synergistic emergence of these peaks in BA.2 may be interpreted as a sign of epistatic interactions in which residues 496-505 (including R498 and H505 sites) can acquire the greater allosteric potential in the presence of the N501Y. In addition, we also noticed that a secondary peak of the distribution is located near residues 446-455.

These findings agree with the illuminating experimental studies showing that strong high-order epistasis with Q498R/N501Y pair could reduce binding affinity cost of immune escaping mutations.^113, 114^ According to our results, epistatic couplings between R498, Y501 and H505 can modulate not only the binding affinity to AC2 but also determine allosteric communications and preferential routes of signal transmission in the Omicron complexes.

Structural mapping of the distribution peaks for BA.1 (Figure 9C) and BA.2 (Figure 9D) illustrates the key differences in allosteric communication ensembles. Although the overall disposition of the mediating positions is similar in these systems, the ensemble of potential communication routes in the BA.2 RBD is strongly dominated by the R498/Y501 binding hub and is likely more robust to perturbations due to the increased RBD stability. Intriguingly, most of the Omicron mutational sites are located in the immediate vicinity of major allosteric control points. In network terms, mutations of predicted allosteric residues could cause network rewiring and may compromise the spike mechanism. Our results suggest that Omicron mutations for all studied variants tend to avoid direct targeting of allosterically sensitive sites to retain the spike activity while leveraging their structural proximity to strategic effector positions in the RBD to modulate communication between distant RBD and ACE2 regions.

A similar picture emerged from the analysis of BA.3 and BA.4/BA.5 complexes (Figure 10). The distribution and structural maps of allosteric routes in BA.3 is similar to BA.2 (Figure 10A,B). However, we observed that BA.4/BA.5 mutations have a distinct modulation profile (Figure 10C) resulting in the sparse and diffuse distribution of mediating centers and potential communication routes (Figure 10D).

These results also reflect the highly dynamic BA.4/BA.5 RBD and are consistent with the PRS analysis suggesting that the RBM region may be the preferential hub for sensing and transmitting allosteric signals to the ACE2 molecule. These findings provide further support to our hypothesis that allosteric communications in BA.4/BA.5 variant may form a broad ensemble passing through RBM region which may lead to the increased transmission of BA.4/BA.5 variants. In this case, we also observed synergistic emergence of R498 and Y501 peaks of the distribution though these peaks are not as dominant as in the BA.2 complex (Figure 10B,D). We argue that modulation of epistatic couplings between R498, Y501 and H505 in BA.4/BA.5 variant may be linked with the increased flexibility of the RBD and corresponding greater adaptability for immune escape. To summarize, perturbation network scanning of allosteric communications suggested that the key Omicron binding affinity hotspots N501Y and Q498R could also function as central mediators of allosteric interactions and epistatic couplings, ultimately exerting control over balancing act of multiple functional tradeoffs including conformational plasticity, binding and allostery that determine fitness of the Omicron variants.

## Conclusions

In this study, we systematically examined conformational dynamics, stability and binding of the Omicron RBD BA.1, BA.2, BA.3 and BA.4/BA.5 RBD complexes with ACE2 using microsecond atomistic MD simulations, in silico mutational scanning of the RBD residues, PRS analysis and network-based mutational profiling of allosteric communications and epistatic interactions. Microsecond MD simulations of the Omicron RBD-ACE2 complexes revealed distinct dynamic patterns in the structurally similar complex conformations. Consistent with the experimental data, our results showed that BA.2 mutations may induce the increased stabilization of the RBD in the complex with ACE2 which may be directly linked with the higher binding affinity than the other variants, while a considerably greater flexibility is an important dynamic signature of the BA.4/BA.5 variants. The mutational scanning maps of binding interactions for BA.2 and BA.3 variants were still quite similar, which is consistent with the structural studies and binding affinity measurements. At the same time, more significant differences were observed for BA.4/BA.5 RBD where functional positions R403, N417, V445, A475, G476, N477, V486 and N487 sites become more tolerant to substitutions, reflecting a more dynamic and conformationally adaptable RBD structure in this Omicron subvariant. Through perturbation-response scanning approach and modeling of allosteric interaction networks we examine the thermodynamic and allosteric factors of RBD-ACE2 binding across BA.1, BA.2, BA.3 and BA.4/BA.5 variants. Our findings suggested that BA.2 mutations may not only induce the increased stabilization of the RBD and enhanced binding interface, but also amplify the allosteric potential of the key binding hotspots, thereby increasing the efficiency of preferential routes for long-range communications with ACE2. Network-based mutational profiling approaches probed the effect of the different Omicron variants on allosteric communications, revealing hidden roles of Omicron mutations as “plastic and evolutionary adaptable” modulators of stability, binding and allostery. Through perturbation network scanning of allosteric residue potentials in the Omicron variant complexes, which is performed in the background of the original strain, we identified that the key Omicron binding affinity hotspots N501Y and Q498R could mediate allosteric interactions and epistatic couplings. These results provided a plausible rationale to the experimentally observed epistatic relationships in which Omicron mutations that reduce ACE2 binding but are important for immune escape are compensated via strong epistatic interactions with the binding affinity hotspots R498 and Y501. The findings support a mechanism according to which Omicron mutations may have evolved to balance thermodynamic stability and conformational adaptability in order to ensure proper tradeoff between stability, binding and immune escape.

## Author Contributions

Conceptualization, G.V. and P.T.; methodology, G.V. and P.T.; software, G.V., S.X., M.A., G.G. and P.T.; validation, G.V.; formal analysis, G.V., M.A., G.G., S.X., and P.T. investigation, G.V. and P.T.; resources, G.V., M.A. and G.G.; data curation, G.V.; writing—original draft preparation, G.V.; writing—review and editing, G.V., M.A. and G.G.; visualization, G.V.; supervision, G.V.; project administration, G.V.; funding acquisition, G.V. All authors have read and agreed to the published version of the manuscript.

## Data Availability Statement

Data is fully contained within the article. Crystal structures were obtained and downloaded from the Protein Data Bank (http://www.rcsb.org). All simulationswere performed using openMM high-performance toolkit for molecular simulation that was obtained from websites https://openmm.org/; https://simtk.org/projects/openmm and https://github.com/openmm/openmm. All simulations were performed using the all-atom additive CHARMM36 protein force field that can be obtained from http://mackerell.umaryland.edu/charmm_ff.shtml. The residue interaction network files were obtained for all structures using the Residue Interaction Network Generator (RING) program RING v2.0.1 freely available at http://old.protein.bio.unipd.it/ring/. The rendering of protein structures was done with interactive visualization program UCSF ChimeraX package (https://www.rbvi.ucsf.edu/chimerax/) and Pymol (https://pymol.org/2/) .

All the data obtained in this work (including simulation trajectories, topology and parameter files, molecular dynamics analysis tools, and the in-house scripts are freely available at DOI:10.5281/zenodo.7889040 (https://zenodo.org/record/7889041#.ZFKRVs7MI2w) and in the GitHub sites https://github.com/smu-tao-group/protein-VAE; https://github.com/smu-tao-group/PASSer2.0

## Acknowledgments

The authors acknowledge support from Schmid College of Science and Technology at Chapman University for providing computing resources at the Keck Center for Science and Engineering.

## References

1. Tai, W.; He, L.; Zhang, X.; Pu, J.; Voronin, D.; Jiang, S.; Zhou, Y.; Du, L. Characterization of the receptor-binding domain (RBD) of 2019 novel coronavirus: implication for development of RBD protein as a viral attachment inhibitor and vaccine. Cell. Mol. Immunol. 2020, 17, 613–620. doi: 10.1038/s41423-020-0400-4.

2. Wang, Q.; Zhang, Y.; Wu, L.; Niu, S.; Song, C.; Zhang, Z.; Lu, G.; Qiao, C.; Hu, Y.; Yuen, K. Y.; Wang, Q.; Zhou, H.; Yan, J.; Qi, J. Structural and functional basis of SARS-CoV-2 entry by using human ACE2. Cell 2020, 181, 894–904.e9. doi: 10.1016/j.cell.2020.03.045.

3. Walls, A. C.; Park, Y. J.; Tortorici, M. A.; Wall, A.; McGuire, A. T.; Veesler, D. Structure, Function, and Antigenicity of the SARS-CoV-2 Spike Glycoprotein. Cell 2020, 181, 281–292.e6. doi: 10.1016/j.cell.2020.02.058.

4. Wrapp, D.; Wang, N.; Corbett, K. S.; Goldsmith, J. A.; Hsieh, C. L.; Abiona, O.; Graham, B. S.; McLellan, J. S. Cryo-EM structure of the 2019-nCoV spike in the prefusion conformation. Science 2020, 367, 1260–1263. doi: 10.1126/science.abb2507.

5. Cai, Y.; Zhang, J.; Xiao, T.; Peng, H.; Sterling, S. M.; Walsh, R. M., Jr.; Rawson, S.; Rits-Volloch, S.; Chen, B. Distinct conformational states of SARS-CoV-2 spike protein. Science 2020, 369, 1586–1592. doi: 10.1126/science.abd4251.

6. Hsieh, C. L.; Goldsmith, J. A.; Schaub, J. M.; DiVenere, A. M.; Kuo, H. C.; Javanmardi, K.; Le, K. C.; Wrapp, D.; Lee, A. G.; Liu, Y., Chou, C.W.; Byrne, P.O.; Hjorth, C.K.; Johnson, N.V.; Ludes-Meyers J.; Nguyen, A.W.; Park, J.; Wang, N.; Amengor, D.; Lavinder, J.J.; Ippolito, G.C.; Maynard, J.A.; Finkelstein, I.J.; McLellan, J.S. Structure-based design of prefusion-stabilized SARS-CoV-2 spikes. Science 2020, 369, 1501–1505. doi: 10.1126/science.abd0826.

7. Henderson, R.; Edwards, R. J.; Mansouri, K.; Janowska, K.; Stalls, V.; Gobeil, S. M. C.; Kopp, M.; Li, D.; Parks, R.; Hsu, A. L., Borgnia, M.J.; Haynes, B.F.; Acharya, P. Controlling the SARS-CoV-2 spike glycoprotein conformation. Nat. Struct. Mol. Biol. 2020, 27, 925–933. doi: 10.1038/s41594-020-0479-4.

8. McCallum, M.; Walls, A. C.; Bowen, J. E.; Corti, D.; Veesler, D. Structure-guided covalent stabilization of coronavirus spike glycoprotein trimers in the closed conformation. Nat. Struct. Mol. Biol. 2020, 27, 942–949. doi: 10.1038/s41594-020-0483-8.

9. Xiong, X.; Qu, K.; Ciazynska, K. A.; Hosmillo, M.; Carter, A. P.; Ebrahimi, S.; Ke, Z.; Scheres, S. H. W.;Bergamaschi, L.; Grice, G. L., Zhang, Y.; CITIID-NIHR COVID-19 BioResource Collaboration, Nathan, J.A.; Baker, S.; James, L.C.; Baxendale, H.E.; Goodfellow, I.; Doffinger, R.; Briggs, J.A.G. A thermostable, closed SARS-CoV-2 spike protein trimer. Nat. Struct. Mol. Biol. 2020, 27, 934–941. doi: 10.1038/s41594-020-0478-5.

10. Costello, S.M.; Shoemaker, S.R.; Hobbs, H.T.; Nguyen, A.W.; Hsieh, C.L.; Maynard, J.A.; McLellan, J.S.; Pak, J.E.; Marqusee, S. The SARS-CoV-2 spike reversibly samples an open-trimer conformation exposing novel epitopes. Nat. Struct. Mol. Biol. 2022, 27, 229–238. doi: 10.1038/s41594-022-00735-5.

11. McCormick, K.D.; Jacobs, J.L.; Mellors, J.W. The emerging plasticity of SARS-CoV-2. Science 2021, 371, 1306–1308. doi: 10.1126/science.abg4493.

12. Ghimire, D.; Han, Y.; Lu, M. Structural Plasticity and Immune Evasion of SARS-CoV-2 Spike Variants. Viruses 2022, 14, 1255. https://doi.org/10.3390/v14061255.

13. Xu, C.; Wang, Y.; Liu, C.; Zhang, C.; Han,W.; Hong, X.; Wang, Y.; Hong, Q.; Wang, S.; Zhao, Q.; Wang, Y.; Yang, Y.; Chen, K.; Zheng, W.; Kong, L.; Wang, F.; Zuo, Q.; Huang, Z.; Cong, Y. Conformational dynamics of SARS-CoV-2 trimeric spike glycoprotein in complex with receptor ACE2 revealed by cryo-EM. Sci. Adv. 2021, 7, eabe5575. doi: 10.1126/sciadv.abe5575.

14. Benton, D. J.; Wrobel, A. G.; Xu, P.; Roustan, C.; Martin, S. R.; Rosenthal, P. B.; Skehel, J. J.; Gamblin, S. J. Receptor binding and priming of the spike protein of SARS-CoV-2 for membrane fusion. Nature 2020, 588, 327–330. doi: 10.1038/s41586-020-2772-0.

15. Turoňová, B.; Sikora, M.; Schürmann, C.; Hagen, W. J. H.; Welsch, S.; Blanc, F. E. C.; von Bülow, S.; Gecht, M.; Bagola, K.; Hörner, C.; van Zandbergen, G.; Landry, J.; de Azevedo, N. T. D.; Mosalaganti, S.; Schwarz, A.; Covino, R.; Mühlebach, M. D.; Hummer, G.; Krijnse Locker, J.; Beck, M. In situ structural analysis of SARS-CoV-2 spike reveals flexibility mediated by three hinges. Science 2020, 370, 203–208. doi: 10.1126/science.abd5223.

16. Lu, M.; Uchil, P. D.; Li, W.; Zheng, D.; Terry, D. S.; Gorman, J.; Shi, W.; Zhang, B.; Zhou, T.; Ding, S.; Gasser, R.; Prevost, J.; Beaudoin-Bussieres, G.; Anand, S. P.; Laumaea, A.; Grover, J. R.; Lihong, L.; Ho, D. D.; Mascola, J.R.; Finzi, A.; Kwong, P. D.; Blanchard, S. C.; Mothes, W. Real-time conformational dynamics of SARS-CoV-2 spikes on virus particles. Cell Host Microbe. 2020, 28, 880–891.e8. doi: 10.1016/j.chom.2020.11.001.

17. Yang, Z.; Han, Y.; Ding, S.; Shi, W.; Zhou, T.; Finzi, A.; Kwong, P.D.; Mothes, W.; Lu, M. SARS-CoV-2 Variants Increase Kinetic Stability of Open Spike Conformations as an Evolutionary Strategy. mBio 2022, 13, e0322721. doi: 10.1128/mbio.03227-21.

18. Díaz-Salinas, M.A.; Li, Q.; Ejemel, M.; Yurkovetskiy, L.; Luban, J.; Shen, K.; Wang, Y.; Munro, J.B. Conformational dynamics and allosteric modulation of the SARS-CoV-2 spike. Elife 2022, 11, e75433. doi: 10.7554/eLife.75433.

19. Han, P.; Li, L.; Liu, S.; Wang, Q.; Zhang, D.; Xu, Z.; Li, X.; Peng, Q.; Su, C.; Huang, B.; Li, D.; Zhang, R.; Tian, M.; Fu, L.; Gao, Y.; Zhao, X.; Liu, K.; Qi, J.; Gao, G. F.; Wang, P. Receptor binding and complex structures of human ACE2 to spike RBD from omicron and delta SARS-CoV-2. Cell 2022, doi: 10.1016/j.cell.2022.01.001.

20. Saville, J.W.; Mannar, D.; Zhu, X.; Srivastava, S.S.; Berezuk, A.M.; Demers, J.P.; Zhou, S.; Tuttle, K.S.; Sekirov, I.; Kim A.; Li, W.; Dimitrov, D.S.; Subramaniam, S. Structural and biochemical rationale for enhanced spike protein fitness in delta and kappa SARS CoV-2 variants. Nat. Commun. 2022, 13, 742. doi: 10.1038/s41467-022-28324-6.

21. Wang, Y.; Liu, C.; Zhang, C.; Wang, Y.; Hong, Q.; Xu, S.; Li, Z.; Yang, Y.; Huang, Z.; Cong, Y. Structural basis for SARS-CoV-2 Delta variant recognition of ACE2 receptor and broadly neutralizing antibodies. Nat. Commun. 2022, 13, 871. doi: 10.1038/s41467-022-28528-w.

22. Zhang, J.; Xiao, T.; Cai, Y.; Lavine, C.L.; Peng, H.; Zhu, H.; Anand, K.; Tong, P.; Gautam, A.; Mayer, M.L.; Walsh, R.M. Jr.; Rits-Volloch, S.; Wesemann, D.R.; Yang, W.; Seaman, M.S.; Lu, J.; Chen, B. Membrane fusion and immune evasion by the spike protein of SARS-CoV-2 Delta variant. Science 2021, 374, 1353–1360. doi: 10.1126/science.abl9463.

23. Mannar, D.; Saville, J.W.; Zhu, X.; Srivastava, S.S.; Berezuk, A.M.; Tuttle, K.S.; Marquez, A.C.; Sekirov, I.; Subramaniam, S. SARS-CoV-2 Omicron variant: Antibody evasion and cryo-EM structure of spike protein-ACE2 complex. Science 2022, 375, 760–764. doi: 10.1126/science.abn7760.

24. Hong, Q.; Han, W.; Li, J.; Xu, S.; Wang, Y.; Xu, C.; Li, Z.; Wang, Y.; Zhang, C.; Huang, Z.; Cong, Y. Molecular basis of receptor binding and antibody neutralization of Omicron. Nature 2022. doi: 10.1038/s41586-022-04581-9.

25. McCallum, M.; Czudnochowski, N.; Rosen, L.E.; Zepeda, S.K.; Bowen, J.E.; Walls, A.C.; Hauser, K.; Joshi, A.; Stewart, C.; Dillen, J.R.; Powell, A.E.; Croll, T.I.; Nix, J.; Virgin, H.W.; Corti, D.; Snell, G.; Veesler, D. Structural basis of SARS-CoV-2 Omicron immune evasion and receptor engagement. Science 2022, 375, 864–868. doi: 10.1126/science.abn8652.

26. Yin, W.; Xu, Y.; Xu, P.; Cao, X.; Wu, C.; Gu, C.; He, X.; Wang, X.; Huang, S.; Wang, X.; Song, B.; Zheng, J.; Jiang, H.; Cheng, X.; Jiang, Y.; Deng, S.J.; Xu, H.E. Structures of the Omicron Spike trimer with ACE2 and an anti-Omicron antibody. Science 2022, 375, 1048–1053. doi: 10.1126/science.abn8863.

27. Gobeil, S.M.; Henderson, R.; Stalls, V.; Janowska, K.; Huang, X.; May, A.; Speakman, M.; Beaudoin, E.; Manne, K.; Li, D.; Parks, R.; Barr, M.; Deyton, M.; Martin, M.; Mansouri, K.; Edwards, R.J.; Sempowski, G.D.; Saunders, K.O.; Wiehe, K.; Williams, W.; Korber, B.; Haynes, B.F.; Acharya, P. Structural diversity of the SARS-CoV 2 Omicron spike. bioRxiv, 2022, doi: 10.1101/2022.01.25.477784.

28. Cui, Z.; Liu, P.; Wang, N.; Wang, L.; Fan, K.; Zhu, Q.; Wang, K.; Chen, R.; Feng, R.; Jia, Z.; Yang, M.; Xu, G.; Zhu, B.; Fu, W.; Chu, T.; Feng, L.; Wang, Y.; Pei, X.; Yang, P.; Xie, X.S.; Cao, L.; Cao, Y.; Wang, X. Structural and functional characterizations of infectivity and immune evasion of SARS-CoV-2 Omicron. Cell 2022, 185, 860–871.e13. doi: 10.1016/j.cell.2022.01.019.

29. Zhou, T.; Wang, L.; Misasi, J.; Pegu, A.; Zhang, Y.; Harris, D.R.; Olia, A.S.; Talana, C.A.; Yang, E.S.; Chen, M.; Choe, M.; Shi, W.; Teng, I.T.; Creanga, A.; Jenkins, C.; Leung, K.; Liu, T.; Stancofski, E.D.; Stephens, T.; Zhang, B.; Tsybovsky, Y.; Graham, B.S.; Mascola, J.R.; Sullivan, N.J.; Kwong, P.D. Structural basis for potent antibody neutralization of SARS-CoV-2 variants including B.1.1.529. Science 2022, 376, eabn8897. doi: 10.1126/science.abn8897.

30. Guo, H.; Gao, Y.; Li, T.; Li, T.; Lu, Y.; Zheng, L.; Liu, Y.; Yang, T.; Luo, F.; Song, S.; Wang, W.; Yang, X.; Nguyen, H. C.; Zhang, H.; Huang, A.; Jin, A.; Yang, H.; Rao, Z.; Ji, X. Structures of Omicron Spike Complexes and Implications for Neutralizing Antibody Development. Cell Rep. 2022, 39, 110770. https://doi.org/10.1016/j.celrep.2022.110770.

31. Stalls, V.; Lindenberger, J.; Gobeil, S. M.-C.; Henderson, R.; Parks, R.; Barr, M.; Deyton, M.; Martin, M.; Janowska, K.; Huang, X.; May, A.; Speakman, M.; Beaudoin, E.; Kraft, B.; Lu, X.; Edwards, R. J.; Eaton, A.; Montefiori, D. C.; Williams, W. B.; Saunders, K. O.; Wiehe, K.; Haynes, B. F.; Acharya, P. Cryo-EM Structures of SARS-CoV-2 Omicron BA.2 Spike. Cell Rep. 2022, 39, 111009. doi: 10.1016/j.celrep.2022.111009.

32. Lin, S.; Chen, Z.; Zhang, X.; Wen, A.; Yuan, X.; Yu, C.; Yang, J.; He, B.; Cao, Y.; Lu, G. Characterization of SARS-CoV-2 Omicron Spike RBD Reveals Significantly Decreased Stability, Severe Evasion of Neutralizing-Antibody Recognition but Unaffected Engagement by Decoy ACE2 Modified for Enhanced RBD Binding. Signal Transduct Target Ther. 2022, 7, 6. doi: 10.1038/s41392-022-00914-2.

33. Zhao, Z.; Zhou, J.; Tian, M.; Huang, M.; Liu, S.; Xie, Y.; Han, P.; Bai, C.; Han, P.; Zheng, A, Fu, L.; Gao, Y.; Peng, Q.; Li, Y.; Chai, Y.; Zhang, Z.; Zhao, X.; Song, H.; Qi, J.; Wang, Q.; Wang, P.; Gao, G. F. Omicron SARS-CoV-2 Mutations Stabilize Spike up-RBD Conformation and Lead to a Non-RBM-Binding Monoclonal Antibody Escape. Nat Commun. 2022, 13, 4958. doi: 10.1038/s41467-022-32665-7.

34. Cerutti, G.; Guo, Y.; Liu, L.; Liu, L.; Zhang, Z.; Luo, Y.; Huang, Y.; Wang, H. H.; Ho, D. D.; Sheng, Z.; Shapiro, L. Cryo-EM Structure of the SARS-CoV-2 Omicron Spike. Cell Rep. 2022, 38, 110428. doi: 10.1016/j.celrep.2022.110428.

35. Ye, G.; Liu, B.; Li, F. Cryo-EM Structure of a SARS-CoV-2 Omicron Spike Protein Ectodomain. Nat Commun. 2022, 13, 1214. doi: 10.1038/s41467-022-28882-9.

36. Dejnirattisai, W.; Huo, J.; Zhou, D.; Zahradník, J.; Supasa, P.; Liu, C.; Duyvesteyn, H. M. E.; Ginn, H. M.; Mentzer, A. J.; Tuekprakhon, A.; Nutalai, R.; Wang, B.; Dijokaite, A.; Khan, S.; Avinoam, O.; Bahar, M.; Skelly, D.; Adele, S.; Johnson, S. A.; Amini, A.; Ritter, T. G.; Mason, C.; Dold, C.; Pan, D.; Assadi, S.; Bellass, A.; Omo-Dare, N.; Koeckerling, D.; Flaxman, A.; Jenkin, D.; Aley, P. K.; Voysey, M.; Costa Clemens, S. A.; Naveca, F. G.; Nascimento, V.; Nascimento, F.; Fernandes da Costa, C.; Resende, P. C.; Pauvolid-Correa, A.; Siqueira, M. M.; Baillie, V.; Serafin, N.; Kwatra, G.; Da Silva, K.; Madhi, S. A.; Nunes, M. C.; Malik, T.; Openshaw, P. J. M.; Baillie, J. K.; Semple, M. G.; Townsend, A. R.; Huang, K. A.; Tan, T. K.; Carroll, M. W.; Klenerman, P.; Barnes, E.; Dunachie, S. J.; Constantinides, B.; Webster, H.; Crook, D.; Pollard, A. J.; Lambe, T.; Paterson, N. G.; Williams, M. A.; Hall, D. R.; Fry, E. E.; Mongkolsapaya, J.; Ren, J.; Schreiber, G.; Stuart, D. I.; Screaton, G. R. SARS-CoV-2 Omicron-B.1.1.529 leads to widespread escape from neutralizing antibody responses. Cell 2022, 185, 467–484.e415. doi: 10.1016/j.cell.2021.12.046.

37. Cameroni, E.; Bowen, J. E.; Rosen, L. E.; Saliba, C.; Zepeda, S. K.; Culap, K.; Pinto, D.; VanBlargan, L. A.; De Marco, A.; di Iulio, J.; Zatta, F.; Kaiser, H.; Noack, J.; Farhat, N.; Czudnochowski, N.; Havenar-Daughton, C.; Sprouse, K. R.; Dillen, J. R.; Powell, A. E.; Chen, A.; Maher, C.; Yin, L.; Sun, D.; Soriaga, L.; Bassi, J.; Silacci-Fregni, C.; Gustafsson, C.; Franko, N. M.; Logue, J.; Iqbal, N. T.; Mazzitelli, I.; Geffner, J.; Grifantini, R.; Chu, H.; Gori, A.; Riva, A.; Giannini, O.; Ceschi, A.; Ferrari, P.; Cippà, P. E.; Franzetti-Pellanda, A.; Garzoni, C.; Halfmann, P. J.; Kawaoka, Y.; Hebner, C.; Purcell, L. A.; Piccoli, L.; Pizzuto, M. S.; Walls, A. C.; Diamond, M. S.; Telenti, A.; Virgin, H. W.; Lanzavecchia, A.; Snell, G.; Veesler, D.; Corti, D. Broadly neutralizing antibodies overcome SARS-CoV-2 Omicron antigenic shift. Nature 2022, 602, 664–670. doi: 10.1038/s41586-021-04386-2.

38. Barton, M.I.; MacGowan, S.A.; Kutuzov, M.A.; Dushek, O.; Barton, G.J.; van der Merwe, P.A. Effects of common mutations in the SARS-CoV-2 Spike RBD and its ligand, the human ACE2 receptor on binding affinity and kinetics. Elife 2021, 10, e70658. doi: 10.7554/eLife.70658.

39. Cao, Y.; Wang, J.; Jian, F.; Xiao, T.; Song, W.; Yisimayi, A.; Huang, W.; Li, Q.; Wang, P.; An, R.; Wang, J.; Wang, Y.; Niu, X.; Yang, S.; Liang, H.; Sun, H.; Li, T.; Yu, Y.; Cui, Q.; Liu, S.; Yang, X.; Du, S.; Zhang, Z.; Hao, X.; Shao, F.; Jin, R.; Wang, X.; Xiao, J.; Wang, Y.; Xie, X.S. Omicron escapes the majority of existing SARS CoV-2 neutralizing antibodies. Nature 2022, 602, 657–663. doi: 10.1038/s41586-021-04385-3.

40. Liu, L.; Iketani, S.; Guo, Y.; Chan, J.F.; Wang, M.; Liu, L.; Luo, Y.; Chu, H.; Huang, Y.; Nair, M.S.; Yu, J.; Chik, K. K.; Yuen, T.T.; Yoon, C.; To, K.K.; Chen, H.; Yin, M.T.; Sobieszczyk, M.E.; Huang, Y.; Wang, H.H.; Sheng, Z.; Yuen, K.Y.; Ho, D.D. Striking antibody evasion manifested by the Omicron variant of SARS-CoV-2. Nature 2022, 602, 676–681. doi: 10.1038/s41586-021-04388-0.

41. Zhang, J.; Cai, Y.; Lavine, C.L.; Peng, H.; Zhu, H.; Anand, K.; Tong, P.; Gautam, A.; Mayer, M.L.; Rits-Volloch, S.; Wang, S.; Sliz, P.; Wesemann, D.R.; Yang, W.; Seaman, M.S.; Lu, J.; Xiao, T.; Chen, B. Structural and functional impact by SARS-CoV-2 Omicron spike mutations. Cell Rep. 2022, 39, 110729. doi: 10.1016/j.celrep.2022.110729.

42. Zhu, R.; Canena, D.; Sikora, M.; Klausberger, M.; Seferovic, H.; Mehdipour, A. R.; Hain, L.; Laurent, E.; Monteil, V.; Wirnsberger, G.; Wieneke, R.; Tampé, R.; Kienzl, N. F.; Mach, L.; Mirazimi, A.; Oh, Y. J.; Penninger, J. M.; Hummer, G.; Hinterdorfer, P. Force-Tuned Avidity of Spike Variant-ACE2 Interactions Viewed on the Single-Molecule Level. Nat Commun. 2022, 13, 7926. doi: 10.1038/s41467-022-35641-3.

43. Bauer, M.S.; Gruber, S.; Hausch, A.; Gomes, P.S.F.C.; Milles, L.F.; Nicolaus, T.; Schendel, L.C.; Navajas, P.L.; Procko, E.; Lietha, D.; Melo, M.C.R.; Bernardi, R.C.; Gaub, H.E.; Lipfert, J. A tethered ligand assay to probe SARS-CoV-2:ACE2 interactions. Proc. Natl. Acad. Sci. U .S. A. 2022, 119, e2114397119. doi: 10.1073/pnas.2114397119. A tethered ligand assay to probe SARS-CoV-2:ACE2 interactions. *Proc. Natl. Acad. Sci. U .S. A.* 2022, 119, e2114397119. D1oi: 10.1073/pnas.2114397119.

44. Hu, W.; Zhang, Y.; Fei, P.; Zhang, T.; Yao, D.; Gao, Y.; Liu, J.; Chen, H.; Lu, Q.; Mudianto, T.; Zhang, X.; Xiao, C.; Ye, Y.; Sun, Q.; Zhang, J.; Xie, Q.; Wang, P.H.; Wang, J.; Li, Z.; Lou, J.; Chen, W. Mechanical activation of spike fosters SARS-CoV-2 viral infection. Cell Res. 2021, 31, 1047–1060. doi: 10.1038/s41422-021-00558-x.

45. Li, L.; Liao, H.; Meng, Y.; Li, W.; Han, P.; Liu, K.; Wang, Q.; Li, D.; Zhang, Y.; Wang, L.; Fan, Z.; Zhang, Y.; Wang, Q.; Zhao, X.; Sun, Y.; Huang, N.; Qi, J.; Gao, G.F. Structural basis of human ACE2 higher binding affinity to currently circulating Omicron SARS-CoV-2 sub-variants BA.2 and BA.1.1. Cell 2022, 185, 2952–2960.e10. doi: 10.1016/j.cell.2022.06.023.

46. Xu, Y.; Wu, C.; Cao, X.; Gu, C.; Liu, H.; Jiang, M.; Wang, X.; Yuan, Q.; Wu, K.; Liu, J.; Wang, D.; He, X.; Wang, X.; Deng, S.J.; Xu, H.E.; Yin, W. Structural and biochemical mechanism for increased infectivity and immune evasion of Omicron BA.2 variant compared to BA.1 and their possible mouse origins. Cell Res. 2022, 32, 609–620. doi: 10.1038/s41422-022-00672-4.

47. Tuekprakhon, A.; Nutalai, R.; Dijokaite-Guraliuc, A.; Zhou, D.; Ginn, H.M.; Selvaraj, M.; Liu, C.; Mentzer, A.J.; Supasa, P.; Duyvesteyn, H.M.E.; Das, R.; Skelly, D.; Ritter, T.G.; Amini, A.; Bibi, S.; Adele, S.; Johnson, S.A.; Constantinides, B.; Webster, H.; Temperton, N.; Klenerman, P.; Barnes, E.; Dunachie, S.J.; Crook, D.; Pollard, A.J.; Lambe, T.; Goulder, P.; Paterson, N.G.; Williams, M.A.; Hall DR; OPTIC Consortium; ISARIC4C Consortium; Fry, E.E.; Huo, J.; Mongkolsapaya, J.; Ren, J.; Stuart, D.I.; Screaton, G.R. Antibody escape of SARS-CoV-2 Omicron BA.4 and BA.5 from vaccine and BA.1 serum. Cell 2022, 185, 2422–2433.e13. doi: 10.1016/j.cell.2022.06.005.

48. Cao, Y.; Yisimayi, A.; Jian, F.; Song, W.; Xiao, T.; Wang, L.; Du, S.; Wang, J.; Li, Q.; Chen, X.; Yu, Y.; Wang, P.; Zhang, Z.; Liu, P.; An, R.; Hao, X.; Wang, Y.; Wang, J.; Feng, R.; Sun, H.; Zhao, L.; Zhang, W.; Zhao, D.; Zheng, J.; Yu, L.; Li, C.; Zhang, N.; Wang, R.; Niu, X.; Yang, S.; Song, X.; Chai, Y.; Hu, Y.; Shi, Y.; Zheng, L.; Li, Z.; Gu, Q.; Shao, F.; Huang, W.; Jin, R.; Shen, Z.; Wang, Y.; Wang, X.; Xiao, J.; Xie, X.S. BA.2.12.1, BA.4 and BA.5 escape antibodies elicited by Omicron infection. Nature 2022, 608, 593–602. doi: 10.1038/s41586-022-04980-y.

49. Bowen, J.E.; Addetia, A.; Dang, H.V.; Stewart, C.; Brown, J.T.; Sharkey, W.K.; Sprouse, K.R.; Walls, A.C.; Mazzitelli, I.G.; Logue, J.K.; Franko, N.M.; Czudnochowski, N.; Powell, A.E.; Dellota, E. Jr.; Ahmed, K.; Ansari, A.S.; Cameroni, E.; Gori, A.; Bandera, A.; Posavad, C.M.; Dan, J.M.; Zhang, Z.; Weiskopf, D.; Sette, A.; Crotty, S.; Iqbal, N.T.; Corti, D.; Geffner, J.; Snell, G.; Grifantini, R.; Chu, H.Y.; Veesler, D. Omicron spike function and neutralizing activity elicited by a comprehensive panel of vaccines. Science 2022, 377, 890–894. doi: 10.1126/science.abq0203.

50. Ni, D.; Turelli, P.; Beckert, B.; Nazarov, S.; Uchikawa, E.; Myasnikov, A.; Pojer, F.; Trono, D.; Stahlberg, H.; Lau, K. Cryo-EM Structures and Binding of Mouse and Human ACE2 to SARS-CoV-2 Variants of Concern Indicate That Mutations Enabling Immune Escape Could Expand Host Range. PLOS Pathogens 2023, 19, e1011206. doi: 10.1371/journal.ppat.1011206.

51. Kimura, I.; Yamasoba, D.; Tamura, T.; Nao, N.; Suzuki, T.; Oda, Y.; Mitoma, S.; Ito, J.; Nasser, H.; Zahradnik, J.; Uriu, K.; Fujita, S.; Kosugi, Y.; Wang, L.; Tsuda, M.; Kishimoto, M.; Ito, H.; Suzuki, R.; Shimizu, R.; Begum, M. M.; Yoshimatsu, K.; Kimura, K. T.; Sasaki, J.; Sasaki-Tabata, K.; Yamamoto, Y.; Nagamoto, T.; Kanamune, J.; Kobiyama, K.; Asakura, H.; Nagashima, M.; Sadamasu, K.; Yoshimura, K.; Shirakawa, K.; Takaori-Kondo, A.; Kuramochi, J.; Schreiber, G.; Ishii, K. J.; Hashiguchi, T.; Ikeda, T.; Saito, A.; Fukuhara, T.; Tanaka, S.; Matsuno, K.; Sato, K. Virological Characteristics of the SARS-CoV-2 Omicron BA.2 Subvariants, Including BA.4 and BA.5. Cell 2022, 185, 3992-4007.e16. doi: 10.1016/j.cell.2022.09.018.

52. Huo, J.; Dijokaite-Guraliuc, A.; Liu, C.; Zhou, D.; Ginn, H. M.; Das, R.; Supasa, P.; Selvaraj, M.; Nutalai, R.; Tuekprakhon, A.; Duyvesteyn, H. M. E.; Mentzer, A. J.; Skelly, D.; Ritter, T. G.; Amini, A.; Bibi, S.; Adele, S.; Johnson, S. A.; Paterson, N. G.; Williams, M. A.; Hall, D. R.; Plowright, M.; Newman, T. A. H.; Hornsby, H.; de Silva, T. I.; Temperton, N.; Klenerman, P.; Barnes, E.; Dunachie, S. J.; Pollard, A. J.; Lambe, T.; Goulder, P.; Fry, E. E.; Mongkolsapaya, J.; Ren, J.; Stuart, D. I.; Screaton, G. R. A Delicate Balance between Antibody Evasion and ACE2 Affinity for Omicron BA.2.75. Cell Rep. 2023, 42, 111903. doi: 10.1016/j.celrep.2022.111903.

53. Park, Y.-J.; Pinto, D.; Walls, A. C.; Liu, Z.; De Marco, A.; Benigni, F.; Zatta, F.; Silacci-Fregni, C.; Bassi, J.; Sprouse, K. R.; Addetia, A.; Bowen, J. E.; Stewart, C.; Giurdanella, M.; Saliba, C.; Guarino, B.; Schmid, M. A.; Franko, N. M.; Logue, J. K.; Dang, H. V.; Hauser, K.; di Iulio, J.; Rivera, W.; Schnell, G.; Rajesh, A.; Zhou, J.; Farhat, N.; Kaiser, H.; Montiel-Ruiz, M.; Noack, J.; Lempp, F. A.; Janer, J.; Abdelnabi, R.; Maes, P.; Ferrari, P.; Ceschi, A.; Giannini, O.; de Melo, G. D.; Kergoat, L.; Bourhy, H.; Neyts, J.; Soriaga, L.; Purcell, L. A.; Snell, G.; Whelan, S. P. J.; Lanzavecchia, A.; Virgin, H. W.; Piccoli, L.; Chu, H. Y.; Pizzuto, M. S.; Corti, D.; Veesler, D. Imprinted Antibody Responses against SARS-CoV-2 Omicron Sublineages. Science, 2022, 378, 619–627. doi: 10.1126/science.adc9127.

54. Hachmann, N. P.; Miller, J.; Collier, A. Y.; Ventura, J. D.; Yu, J.; Rowe, M.; Bondzie, E. A.; Powers, O.; Surve, N.; Hall, K.; Barouch, D. H. Neutralization Escape by SARS-CoV-2 Omicron Subvariants BA.2.12.1, BA.4, and BA.5. N. Engl. J. Med. 2022, 387, 86–88. doi: 10.1056/NEJMc2206576.

55. Zhang, W.; Shi, K.; Geng, Q.; Ye, G.; Aihara, H.; Li, F. Structural Basis for Mouse Receptor Recognition by SARS-CoV-2 Omicron Variant. Proc. Natl. Acad. Sci. U .S. A. 2022, 119,e2206509119. doi: 10.1073/pnas.2206509119.

56. Casalino, L.; Gaieb, Z.; Goldsmith, J. A.; Hjorth, C. K.; Dommer, A. C.; Harbison, A. M.; Fogarty, C. A.; Barros, E. P.; Taylor, B. C.; McLellan, J. S.; Fadda, E.; Amaro, R. E., Beyond Shielding: The Roles of Glycans in the SARS-CoV-2 Spike Protein. ACS Cent. Sci. 2020, 6, 1722–1734. doi: 10.1021/acscentsci.0c01056.

57. Sztain, T.; Ahn, S.H.; Bogetti, A.T.; Casalino, L.; Goldsmith, J.A.; Seitz, E.; McCool, R.S.; Kearns, F.L.; Acosta-Reyes, F.; Maji, S.; Mashayekhi, G.; McCammon, J.A.; Ourmazd, A.; Frank, J.; McLellan, J.S.; Chong, L.T.; Amaro, R.E. A glycan gate controls the opening of the SARS-CoV-2 spike protein. Nat. Chem. 2021, doi: 10.1038/s41557-021-00758-3.

58. Sikora, M.; von Bülow, S.; Blanc, F. E. C.; Gecht, M.; Covino, R.; Hummer, G., Computational epitope map of SARS-CoV-2 spike protein. PLoS Comput. Biol. 2021, 17, e1008790. doi: 10.1371/journal.pcbi.1008790.

59. Pang, Y. T.; Acharya, A.; Lynch, D. L.; Pavlova, A.; Gumbart, J. C. SARS-CoV-2 Spike Opening Dynamics and Energetics Reveal the Individual Roles of Glycans and Their Collective Impact. *Commun*. Biol. 2022, 5, 1170. doi: 10.1038/s42003-022-04138-6.

60. Xu, C.; Wang, Y.; Liu, C.; Zhang, C.; Han,W.; Hong, X.; Wang, Y.; Hong, Q.; Wang, S.; Zhao, Q.; Wang, Y.; Yang, Y.; Chen, K.; Zheng, W.; Kong, L.; Wang, F.; Zuo, Q.; Huang, Z.; Cong, Y. Conformational dynamics of SARS-CoV-2 trimeric spike glycoprotein in complex with receptor ACE2 revealed by cryo-EM. Sci. Adv. 2021, 7, eabe5575. doi: 10.1126/sciadv.abe5575.

61. Mori, T.; Jung, J.; Kobayashi, C.; Dokainish, H.M.; Re, S.; Sugita, Y. Elucidation of interactions regulating conformational stability and dynamics of SARS-CoV-2 S-protein. Biophys. J. 2021, 120, 1060–1071. doi: 10.1016/j.bpj.2021.01.012.

62. Zimmerman, M. I.; Porter, J. R.; Ward, M. D.; Singh, S.; Vithani, N.; Meller, A.; Mallimadugula, U. L.; Kuhn, C. E.; Borowsky, J. H.; Wiewiora, R. P., Hurley, M.F.D.; Harbison, A.M.; Fogarty, C.A.; Coffland, J.E.; Fadda, E.; Voelz, V.A.; Chodera, J.D.; Bowman, G.R. SARS-CoV-2 simulations go exascale to predict dramatic spike opening and cryptic pockets across the proteome. Nat. Chem. 2021, 13, 651–659. doi: 10.1038/s41557-021-00707-0.

63. Mori, T.; Jung, J.; Kobayashi, C.; Dokainish, H. M.; Re, S.; Sugita, Y. Elucidation of Interactions Regulating Conformational Stability and Dynamics of SARS-CoV-2 S-Protein. Biophys J. 2021, 120, 1060–1071. doi: 10.1016/j.bpj.2021.01.012.

64. Dokainish, H.M.; Re, S.; Mori, T.; Kobayashi, C.; Jung, J.; Sugita, Y. The inherent flexibility of receptor binding domains in SARS-CoV-2 spike protein. Elife 2022, 11, e75720. doi: 10.7554/eLife.75720.

65. Dokainish, H. M.; Sugita, Y. Structural Effects of Spike Protein D614G Mutation in SARS CoV-2. Biophys J. 2022 Nov 17:S0006–3495(22)00941-9. doi: 10.1016/j.bpj.2022.11.025.

66. Verkhivker, G.M. Coevolution, dynamics and allostery conspire in shaping cooperative binding and signal transmission of the SARS-CoV-2 spike protein with human angiotensin converting enzyme 2. Int. J. Mol. Sci. 2020, 21, 8268. doi: 10.3390/ijms21218268.

67. Verkhivker, G.M. Molecular simulations and network modeling reveal an allosteric signaling in the SARS-CoV-2 spike proteins. J. Proteome Res. 2020, 19, 4587–4608. doi: 10.1021/acs.jproteome.0c00654.

68. Verkhivker, G. M.; Di Paola, L.; Dynamic Network Modeling of Allosteric Interactions and Communication Pathways in the SARS-CoV-2 Spike Trimer Mutants: Differential Modulation of Conformational Landscapes and Signal Transmission via Cascades of Regulatory Switches. J Phys. Chem. B. 2021, 125, 850–873. doi: 10.1021/acs.jpcb.0c10637.

69. Verkhivker, G.M.; Di Paola, L. Integrated Biophysical Modeling of the SARS-CoV-2 Spike Protein Binding and Allosteric Interactions with Antibodies. J. Phys. Chem. B. 2021, 125, 4596–4619. doi: 10.1021/acs.jpcb.1c00395.

70. Verkhivker, G.M.; Agajanian, S.; Oztas, D.Y.; Gupta, G. Comparative Perturbation-Based Modeling of the SARS-CoV-2 Spike Protein Binding with Host Receptor and Neutralizing Antibodies: Structurally Adaptable Allosteric Communication Hotspots Define Spike Sites Targeted by Global Circulating Mutations. Biochemistry 2021, 60, 1459–1484. doi: 10.1021/acs.biochem.1c00139.

71. Verkhivker, G.M.; Agajanian, S.; Oztas, D.Y.; Gupta, G. Dynamic Profiling of Binding and Allosteric Propensities of the SARS-CoV-2 Spike Protein with Different Classes of Antibodies: Mutational and Perturbation-Based Scanning Reveals the Allosteric Duality of Functionally Adaptable Hotspots. J. Chem. Theory Comput. 2021, 17, 4578–4598. doi: 10.1021/acs.jctc.1c00372.

72. Verkhivker, G.M.; Agajanian, S.; Oztas, D.Y.; Gupta, G. Allosteric Control of Structural Mimicry and Mutational Escape in the SARS-CoV-2 Spike Protein Complexes with the ACE2 Decoys and Miniprotein Inhibitors: A Network-Based Approach for Mutational Profiling of Binding and Signaling. J. Chem. Inf. Model. 2021, 61, 5172–5191. doi: 10.1021/acs.jcim.1c00766.

73. Hossen, M.L.; Baral, P.; Sharma, T.; Gerstman, B.; Chapagain, P. Significance of the RBD mutations in the SARS-CoV-2 omicron: from spike opening to antibody escape and cell attachment. Phys. Chem. Chem. Phys. 2022, 24, 9123–9129. doi: 10.1039/d2cp00169a.

74. Jawad, B.; Adhikari, P.; Podgornik, R.; Ching, W.Y. Binding Interactions between Receptor-Binding Domain of Spike Protein and Human Angiotensin Converting Enzyme-2 in Omicron Variant. J. Phys. Chem. Lett. 2022, 13, 3915–3921. doi: 10.1021/acs.jpclett.2c00423.

75. Gan, H.H. Twaddle, A.; Marchand, B.; Gunsalus, K.C. Structural Modeling of the SARS-CoV-2 Spike/Human ACE2 Complex Interface can Identify High-Affinity Variants Associated with Increased Transmissibility. J. Mol. Biol. 2021, 433, 167051. doi: 10.1016/j.jmb.2021.167051.

76. Verkhivker, G.; Agajanian, S.; Kassab, R.; Krishnan, K. Computer Simulations and Network-Based Profiling of Binding and Allosteric Interactions of SARS-CoV-2 Spike Variant Complexes and the Host Receptor: Dissecting the Mechanistic Effects of the Delta and Omicron Mutations. Int. J. Mol. Sci. 2022, 23, 4376. doi: 10.3390/ijms23084376.

77. Verkhivker, G.; Agajanian, S.; Kassab, R.; Krishnan, K. Probing Mechanisms of Binding and Allostery in the SARS-CoV-2 Spike Omicron Variant Complexes with the Host Receptor: Revealing Functional Roles of the Binding Hotspots in Mediating Epistatic Effects and Communication with Allosteric Pockets. Int. J. Mol. Sci. 2022, 23, 11542. doi: 10.3390/ijms231911542.

78. Rose, P. W.; Prlic, A.; Altunkaya, A.; Bi, C.; Bradley, A. R.; Christie, C. H.; Costanzo, L. D.; Duarte, J. M.; Dutta, S.; Feng, Z.; Green, R. K.; Goodsell, D. S.; Hudson, B.; Kalro, T.; Lowe, R.; Peisach, E.; Randle, C.; Rose, A. S.; Shao, C.; Tao, Y. P.; Valasatava, Y.; Voigt, M.; Westbrook, J. D.; Woo, J.; Yang, H.; Young, J. Y.; Zardecki, C.; Berman, H. M.; Burley, S. K. The RCSB protein data bank: integrative view of protein, gene and 3D structural information. Nucleic Acids Res. 2017, 45, D271–D281. doi: 10.1093/nar/gkw1000.

79. Hekkelman, M.L.; Te Beek, T.A.; Pettifer, S.R.; Thorne, D.; Attwood, T.K.; Vriend, G. WIWS: A protein structure bioinformatics web service collection. Nucleic Acids Res. 2010, 38, W719–W723. https://doi.org/10.1093/nar/gkq453.

80. Fernandez-Fuentes, N.; Zhai, J.; Fiser, A. ArchPRED: A template based loop structure prediction server. Nucleic Acids Res. 2006, 34, W173–W176. https://doi.org/10.1093/nar/gkl113.

81. Krivov, G.G.; Shapovalov, M.V.; Dunbrack, R.L., Jr. Improved prediction of protein side chain conformations with SCWRL4. Proteins 2009, 77, 778–795. https://doi.org/10.1002/prot.22488.

82. Bhattacharya, D.; Cheng, J. 3Drefine: Consistent Protein Structure Refinement by Optimizing Hydrogen Bonding Network and Atomic-Level Energy Minimization. Proteins 2013, 81, 119–131. doi: 10.1002/prot.24167.

83. Bhattacharya, D.; Nowotny, J.; Cao, R.; Cheng, J. 3Drefine: An Interactive Web Server for Efficient Protein Structure Refinement. Nucleic Acids Res. 2016, 44, W406–W409. doi: 10.1093/nar/gkw336.

84. Huang, J.; Rauscher, S.; Nawrocki, G.; Ran, T.; Feig, M.; de Groot, B.L.; Grubmüller, H.; MacKerell, A.D. Jr. CHARMM36m: an improved force field for folded and intrinsically disordered proteins. Nat Methods 2017, 14, 71–73. doi: 10.1038/nmeth.4067.

85. Jorgensen, W. L.; Chandrasekhar, J.; Madura, J. D.; Impey, R. W.; Klein, M. L. Comparison of Simple Potential Functions for Simulating Liquid Water. J. Chem. Phys. 1983, 79, 926–935. https://doi.org/10.1063/1.445869.

86. Fernandes, H.S.; Sousa, S.F.; Cerqueira, N.M.F.S.A. VMD Store-A VMD Plugin to Browse, Discover, and Install VMD Extensions. J. Chem. Inf. Model. 2019, 59, 4519–4523. doi: 10.1021/acs.jcim.9b00739.

87. Ryckaert, J.-P.; Ciccotti, G.; Berendsen, H. J. C. Numerical Integration of the Cartesian Equations of Motion of a System with Constraints: Molecular Dynamics of n-Alkanes. J. Comput. Phys. 1977, 23, 327–341. https://doi.org/10.1016/0021-9991(77)90098-5.

88. Di Pierro, M.; Elber, R.; Leimkuhler, B. A Stochastic Algorithm for the Isobaric-Isothermal Ensemble with Ewald Summations for All Long Range Forces. J. Chem. Theory Comput. 2015, 11, 5624–5637. https://doi.org/10.1021/acs.jctc.5b00648.

89. Eastman, P.; Swails, J.; Chodera, J. D.; McGibbon, R. T.; Zhao, Y.; Beauchamp, K. A.; Wang, L.-P.; Simmonett, A. C.; Harrigan, M. P.; Stern, C. D.; Wiewiora, R. P.; Brooks, B. R.; Pande, V. S. OpenMM 7: Rapid Development of High Performance Algorithms for Molecular Dynamics. PLoS Comput Biol. 2017, 13, e1005659. doi: 10.1371/journal.pcbi.1005659.

90. Dehouck, Y.; Kwasigroch, J. M.; Rooman, M.; Gilis, D. BeAtMuSiC: Prediction of changes in protein-protein binding affinity on mutations. Nucleic Acids Res. 2013, 41, W333–W339. doi: 10.1093/nar/gkt450.

91. Dehouck, Y.; Gilis, D.; Rooman, M. A new generation of statistical potentials for proteins. Biophys. J. 2006, 90, 4010–4017. doi: 10.1529/biophysj.105.079434.

92. Dehouck, Y.; Grosfils, A.; Folch, B.; Gilis, D.; Bogaerts, P.; Rooman, M. Fast and accurate predictions of protein stability changes upon mutations using statistical potentials and neural networks: PoPMuSiC-2.0. Bioinformatics 2009, 25, 2537–2543. doi: 10.1093/bioinformatics/btp445.

93. Atilgan, C.; Atilgan, A. R., Perturbation-response scanning reveals ligand entry-exit mechanisms of ferric binding protein. PLoS Comput. Biol. 2009, 5, e1000544.

94. Atilgan, C.; Gerek, Z. N.; Ozkan, S. B.; Atilgan, A. R., Manipulation of conformational change in proteins by single-residue perturbations. Biophys. J. 2010, 99, 933–943.

95. Jalalypour, F.; Sensoy, O.; Atilgan, C., Perturb-Scan-Pull: A Novel Method Facilitating Conformational Transitions in Proteins. J. Chem. Theory Comput. 2020, 16, 3825–3841.

96. General, I. J.; Liu, Y.; Blackburn, M. E.; Mao, W.; Gierasch, L. M.; Bahar, I., ATPase subdomain IA is a mediator of interdomain allostery in Hsp70 molecular chaperones. PLoS Comput. Biol. 2014, 10, e1003624.

97. Dutta, A.; Krieger, J.; Lee, J. Y.; Garcia-Nafria, J.; Greger, I. H.; Bahar, I., Cooperative Dynamics of Intact AMPA and NMDA Glutamate Receptors: Similarities and Subfamily Specific Differences. Structure 2015, 23, 1692–1704.

98. Stetz, G.; Tse, A.; Verkhivker, G. M., Dissecting Structure-Encoded Determinants of Allosteric Cross-Talk between Post-Translational Modification Sites in the Hsp90 Chaperones. Sci. Rep. 2018, 8, 19.

99. Verkhivker, G. M.; Di Paola, L., Dynamic Network Modeling of Allosteric Interactions and Communication Pathways in the SARS-CoV-2 Spike Trimer Mutants: Differential Modulation of Conformational Landscapes and Signal Transmission via Cascades of Regulatory Switches. J Phys Chem B 2021, 125, 850–873.

100. Brinda, K.V., Vishveshwara, S. A network representation of protein structures: Implications for protein stability. Biophys. J. 2005, 89, 4159–4170.

101. Vijayabaskar, M.S., and Vishveshwara, S. Interaction energy based protein structure networks. Biophys. J. 2010, 99, 3704–3715.

102. Sethi, A., Eargle, J., Black, A.A., Luthey-Schulten, Z. Dynamical networks in tRNA:protein complexes. Proc. Natl. Acad. Sci. U.S.A. 2009, 106, 6620–6625.

103. Stetz, G., and Verkhivker, G. M. Computational analysis of residue interaction networks and coevolutionary relationships in the Hsp70 chaperones: A community-hopping model of allosteric regulation and communication. Plos Comput. Biol. 2017, 13, e1005299.

104. Martin, A. J. M.; Vidotto, M.; Boscariol, F.; Di Domenico, T.; Walsh, I.; Tosatto, S. C. E. RING: Networking Interacting Residues, Evolutionary Information and Energetics in Protein Structures. Bioinformatics 2011, 27, 2003–2005. doi: 10.1093/bioinformatics/btr191.

105. Del Conte, A.; Monzon, A. M.; Clementel, D.; Camagni, G. F.; Minervini, G.; Tosatto, S. C. E.; Piovesan, D. RING-PyMOL: Residue Interaction Networks of Structural Ensembles and Molecular Dynamics. Bioinformatics 2023. doi: 10.1093/bioinformatics/btad260.

106. Clementel, D.; Del Conte, A.; Monzon, A. M.; Camagni, G. F.; Minervini, G.; Piovesan, D.; Tosatto, S. C. E. RING 3.0: Fast Generation of Probabilistic Residue Interaction Networks from Structural Ensembles. Nucleic Acids Res. 2022, 50, W651–W656. doi: 10.1093/nar/gkac365.

107. Hagberg, A.A.; Schult, D.A.; Swart, P.J. Exploring network structure, dynamics, and function using NetworkX. In Proceedings of the 7th Python in Science Conference (SciPy2008), Pasadena, CA, USA, 19–24 August 2008, Varoquaux, G., Vaught, T., Millman, J., Eds.; Scientific Research: Atlanta, GA, USA, 2011; pp. 11–15.

108. del Sol, A.; Fujihashi, H.; Amoros, D.; Nussinov, R. Residues crucial for maintaining short paths in network communication mediate signaling in proteins. Mol. Syst. Biol. 2006, 2, 2006.0019. doi: 10.1038/msb4100063.

109. Brysbaert, G.; Mauri, T.; Lensink, M. F. Comparing protein structures with RINspector automation in Cytoscape. F1000Res. 2018, 7, 563. doi: 10.12688/f1000research.14298.2.

110. Rössler, A.; Netzl, A.; Knabl, L.; Schäfer, H.; Wilks, S. H.; Bante, D.; Falkensammer, B.; Borena, W.; von Laer, D.; Smith, D. J.; Kimpel, J. BA.2 and BA.5 Omicron Differ Immunologically from Both BA.1 Omicron and Pre-Omicron Variants. Nat Commun. 2022, 13, 7701. doi: 10.1038/s41467-022-35312-3.

111. Starr, T. N.; Greaney, A. J.; Hilton, S. K.; Ellis, D.; Crawford, K. H. D.; Dingens, A. S.; Navarro, M. J.; Bowen, J. E.; Tortorici, M. A.; Walls, A. C.; King, N. P.; Veesler, D.; Bloom, J. D. Deep Mutational Scanning of SARS-CoV-2 Receptor Binding Domain Reveals Constraints on Folding and ACE2 Binding. Cell 2020, 182, 1295–1310.e20. doi: 10.1016/j.cell.2020.08.012.

112. Starr, T. N.; Greaney, A. J.; Stewart, C. M.; Walls, A. C.; Hannon, W. W.; Veesler, D.; Bloom, J. D. Deep Mutational Scans for ACE2 Binding, RBD Expression, and Antibody Escape in the SARS-CoV-2 Omicron BA.1 and BA.2 Receptor-Binding Domains. PLoS Pathog. 2022, 18, e1010951. doi: 10.1371/journal.ppat.1010951.

113. Starr, T.N.; Greaney, A.J.; Hannon, W.W.; Loes, A.N.; Hauser, K.; Dillen, J.R.; Ferri, E.; Farrell, A.G.; Dadonaite, B.; McCallum, M.; Matreyek, K.A.; Corti, D.; Veesler, D.; Snell, G.; Bloom, J.D. Shifting mutational constraints in the SARS-CoV-2 receptor-binding domain during viral evolution. Science 2022, 377, 420–424. doi: 10.1126/science.abo7896.

114. Moulana, A.; Dupic, T.; Phillips, A. M.; Chang, J.; Nieves, S.; Roffler, A. A.; Greaney, A. J.; Starr, T. N.; Bloom, J. D.; Desai, M. M. Compensatory Epistasis Maintains ACE2 Affinity in SARS-CoV-2 Omicron BA.1. Nat Commun. 2022, 13, 7011. doi: 10.1038/s41467-022-34506-z.

