## Supplemental Figures S1,S2 and Tables S1,S2 for "Probing Conformational Landscapes of Binding and Allostery in the SARS-CoV-2 Omicron Variant Complexes Using Microsecond Atomistic Simulations and Perturbation-Based Profiling Approaches: Hidden Role of Omicron Mutations as Modulators of Allosteric Signaling and Epistatic Relationships"

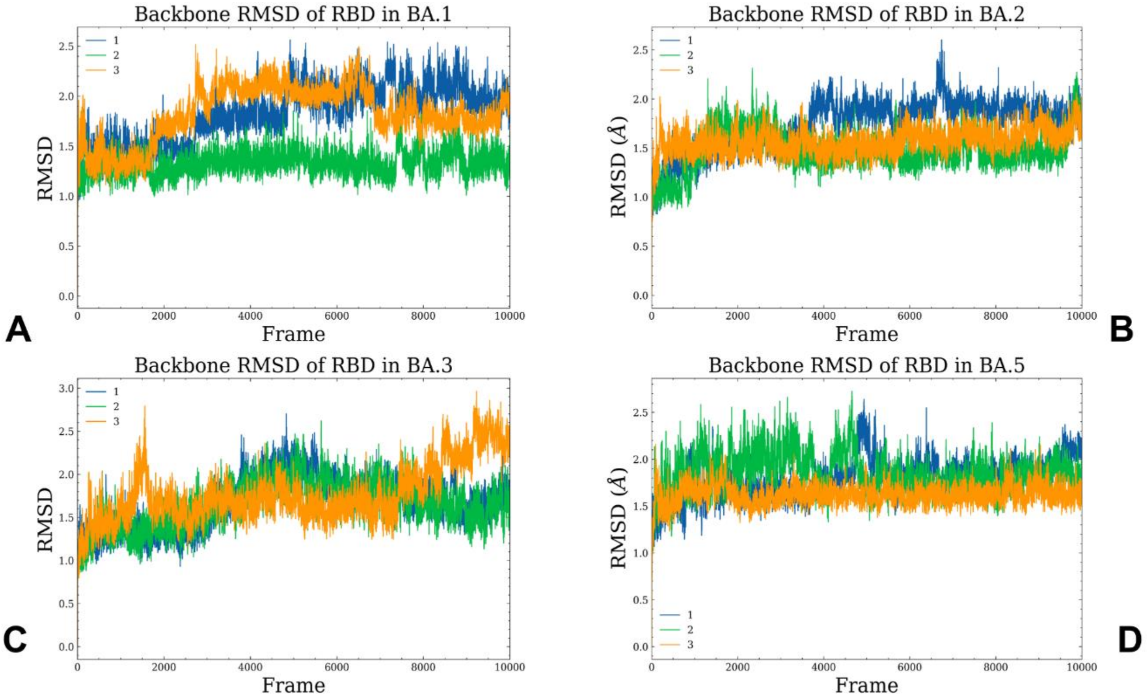


**Figure S1.** Conformational dynamics profiles obtained from all-atom MD simulations of the Omicron RBD BA.1, BA.2, BA.3 and BA.4/BA.5 complexes with hACE2. The RMSD profiles for the RBD residues obtained from 3 microsecond MD simulations of the Omicron RBD BA.1-hACE2 complex, pdb id 7WBP (A), Omicron RBD BA.2-hACE2 complex, pdb id 7XB0 (B), Omicron RBD BA.3-hACE2 complex, pdb id 7XB1 (C) and Omicron RBD BA.4/BA.5-hACE2 complex, pdb id 7XWA (D).

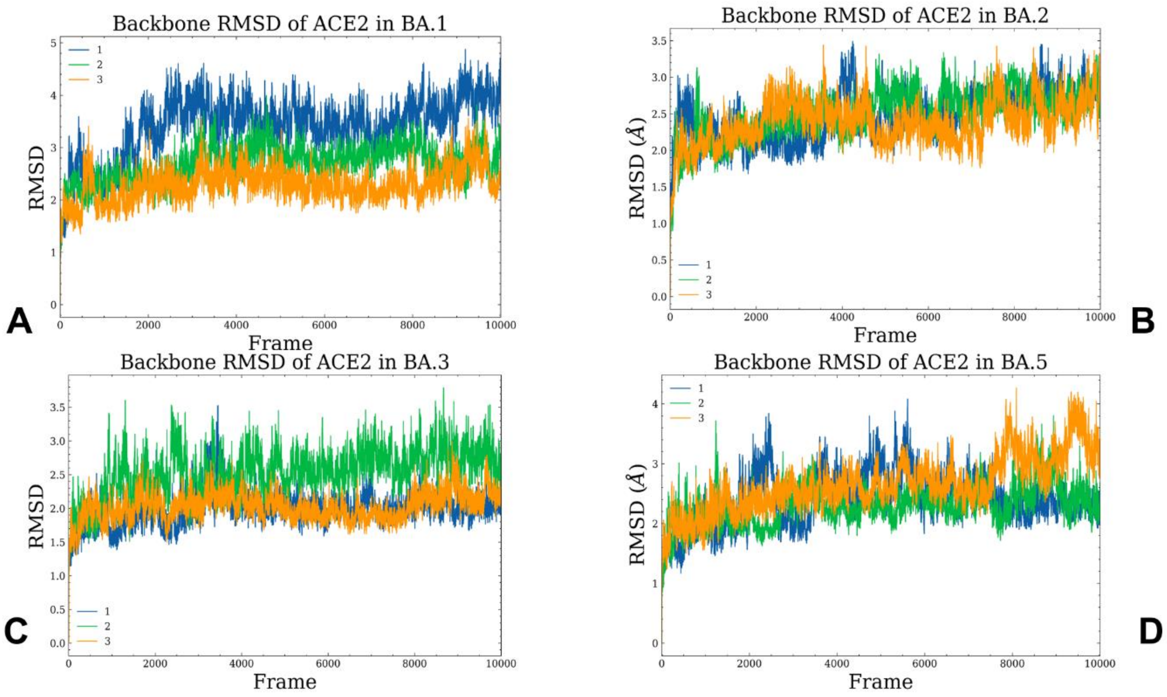


**Figure S2.** Conformational dynamics profiles of the ACE2 residues obtained from MD simulations of the Omicron RBD BA.1, BA.2, BA.3 and BA.4/BA.5 complexes with hACE2. The RMSD profiles for the ACE2 residues obtained from 3 microsecond MD simulations of the Omicron RBD BA.1-hACE2 complex, pdb id 7WBP (A), Omicron RBD BA.2-hACE2 complex, pdb id 7XB0 (B), Omicron RBD BA.3-hACE2 complex, pdb id 7XB1 (C) and Omicron RBD BA.4/BA.5-hACE2 complex, pdb id 7XWA (D).

**Table S1.** Statistical analysis of the intermolecular contact residues in Omicron RBD-hACE2 complexes.*

| ACE2 | BA.1 RBD | BA.2 RBD | BA.3 RBD | BA.4/5 RBD |
|---|---|---|---|---|
| S19 | A475, G476,N477 | A475, G476,N477 | A475, G476,N477 | A475, G476,N477 |
| T20 | | N477 | A475,N477 | N477 |
| Q24 | A475, G476,N477 F486, N487, Y489 | A475, G476,N477 F486,N487, Y489 | A475, G476,N477 F486,N487, Y489 | A475, G476,N477, N487, Y489 |
| T27 | F456, Y473, A475,Y489 | F456, Y473, A475,Y489 | F456, Y473, A475,Y489 | F456,Y473, A475,Y489 |
| F28 | N487,Y489 | N487,Y489 | Y489 | F456, N487,Y489 |
| F30 | L455, F456 | N417, L455, F456 | L455, F456 | L455, F456, Q493 |
| K31 | L455, F456, Y489,R493 | L455, F456, G485,Y489, R493 | L455, F456, Y489,R493 | L455, F456, Y489,Q493 |
| H34 | Y453, L455, R493, S494, Y495 | R403, N417, Y453, L455, R493 | N417, Y453, L455, R493 | N417, Y453, L455, R493 |
| E35 | R493 | R493 | R493 | Q493 |
| E37 | H505 | H505 | H505 | H505 |
| D38 | Y449, S496, R498, Y501 | Y449, Y495, G496, R498, Y501 | Y449, Y495, R498, Y501 | Y449, Y495, G496, R498, Y501 |
| Y41 | R498, T500, Y501 | R498, T500, Y501 | R498, T500, Y501 | R498, T500, Y501 |
| Q42 | S446, Y449, R498 | Y449, R498 | Y449, R498 | Y449, R498 |
| L45 | R498,T500 | V445, R498,T500 | V445, R498,T500 | V445, R498,T500 |
| L79 | F486 | G485, F486 | F486 | V486 |
| M82 | F486 | F486 | F486 | V486 |

| | | | | |
|---|---|---|---|---|
| Y83 | F486, N487, Y489 | F486, N487, Y489 | F486, N487, Y489 | N487, Y489 |
| Q325 | V593 | Q506 | V503, Q506 | |
| G326 | | | T500 | T500 |
| N330 | T500 | P499,T500 | P499,T500 | P499,T500 |
| G352 | | Y501,G502 | Y501,G502 | Y501 |
| K353 | R403, Y495, S496, T500, Y501, G502, H505 | R403,Y495, T500,Y501, G502,V503, H505 | R403, Y495, T500,Y501, G502,V503, H505 | Y495, T500,Y501, G502, H505 |
| G354 | T500,Y501, G502,V503, H505 | T500,Y501, G502,V503, H505 | T500,Y501, G502,V503, H505 | Y501, G502,V503, H505 |
| D355 | T500, Y501,G502 | T500, Y501,G502 | T500, Y501,G502 | T500, Y501,G502 |
| R357 | | T500 | | T500 |

*Two residues are defined in contact if any of their heavy atom is within a distance of 5.0 Å

**Table S2.** The Occupancy of the Pairwise Interactions in the Omicron RBD-hACE2 Complexes

| | Interaction | BA.1-ACE2 | BA.2-ACE2 | BA.3-ACE2 | BA.4/BA.5-ACE2 |
|---|---|---|---|---|---|
| Salt bridges | R403-E37 | 65% | 73% | 73% | 62% |
| | K440-E329 | 31% | 54% | 54% | 53% |
| | R493-E35 | 77% | 92% | 99% | |
| | R493-D38 | 26% | 89% | 89% | |
| | R498-D38 | 59% | 95% | 83% | 78% |
| Hydrophobic Interactions | F456-T27 | 95% | 96% | 88% | 57% |
| | Y473-T27 | 92% | 89% | 85% | 72% |
| | A475-T27 | 88% | 93% | 83% | 66% |
| | F486-F28 | 78% | 97% | 90% | 54% |
| | F486/V486-L79 | 85% | 89% | 82% | 57% |
| | F486/V486-M82 | 85% | 96% | 90% | 62% |
| | F486/V486-Y83 | 90% | 95% | 87% | 53% |
| | Y489-F28 | 97% | 94% | 95% | 86% |
| | Y489-L79 | 90% | 95% | 86% | 72% |
| | Y489-Y83 | 96% | 82% | 88% | 77% |
| Hydrogen Bonds | Y453-H34 | 66% | 82% | 92% | 52% |
| | Y449-D38 | 65% | 82% | 58% | 60% |
| | A475-S19 | 60% | 95% | 85% | 82% |
| | N477-S19 | 58% | 97% | 97% | 69% |
| | N487-Q24 | 62% | 92% | 92% | 71% |
| | N487-Y83 | 76% | 92% | 90% | 54% |
| | Y489-F28 | 82% | 86% | 80% | 68% |
| | T500-D355 | 82% | 77% | 90% | 72% |
| | T500-Y41 | 72% | 80% | 95% | 54% |
| | G502-K353 | 78% | 84% | 78% | 67% |
| | Y501-K353 | 86% | 90% | 84% | 77% |
| | Q493-K31 | | | | 87% |
| | Q493-H34 | | | | 90% |
